# Understanding and addressing under-sampling in plant-pollinator networks using stacked models for missing link prediction

**DOI:** 10.64898/2026.01.08.698368

**Authors:** Lucy Van Kleunen, Natasha de Manincor, Laura E. Dee, Aaron Clauset, Stuart Roberts, François Massol

## Abstract

Pollinator diversity and pollination are under threat from anthropogenic disturbances. Plant-pollinator networks are useful tools to study the consequences of these disturbances, but their links are often under-sampled, potentially biasing conclusions. We introduce a computational method that can be used to predict which unobserved links in a plant-pollinator network dataset are missing. This method extends a state-of-the-art machine learning approach to missing link prediction based on stacked generalization, which has been promising in other contexts, to the setting of bipartite plant-pollinator networks with species traits treated as node attributes. This approach predicts missing links using an ensemble of predictors based on both observed network topology and species traits. We demonstrate this approach on synthetic and empirical plant-pollinator networks. We take advantage of a unique empirical plant-pollinator dataset which samples from three sites using (a) records of observed visits of pollinators to plant species and (b) pollen analysis with detailed species trait annotations for both plant and pollinator nodes. Visit-based sampling under-samples the links in these networks, whereas pollen-based sampling identifies additional links, allowing us to investigate missing link prediction from realistic patterns of field under-sampling. We show that full plant-pollinator networks can be partially reconstructed from the under-sampled networks using our approach. Further, we show that this method can be used to identify which features are the most important for predicting missing links, which allows for investigation into mechanisms driving plant-pollinator link formation and observation. This method is broadly applicable to under-sampled plant-pollinator networks.

## Introduction

Pollinator diversity is under threat from anthropogenic disturbances such as land-use change and the introduction of non-native species (Gill et al., 2016; Potts, 2016; Sánchez-Bayo & Wyckhuys, 2019; Traveset & Richardson, 2014; Vanbergen & Initiative, 2013). Loss of pollinators threatens pollination, as crops that depend on animal pollination contribute to 35% of global crop production value (Potts, 2016); more generally, between 78% to 94% of angiosperms are pollinated by animals (Ollerton et al., 2011). To study the consequences of these changes, plant-pollinator networks provide a useful tool by representing mutualistic pollination interactions in an ecosystem (Ballantyne et al., 2015; Fontaine et al., 2005; Popic et al., 2013). For instance, the structure of networks of mutualistic interactions is fundamental to ecosystem stability (Okuyama & Holland, 2008; Thébault & Fontaine, 2010), and thus plant-pollinator networks have been used to predict the impacts of disturbances on biodiversity when species depend on one another (Valdovinos et al., 2018). Plant-pollinator networks can also be used to understand ecosystem service stability under disturbances and species losses (Keyes et al., 2021; Ross et al., 2021; Valdovinos, 2022).

One challenge for using plant-pollinator networks for forecasting biodiversity and ecosystem service outcomes, however, is that sampling of plant-pollinator networks, like any ecological network, is labor-intensive and costly. As a result, the links in plant-pollinator network datasets are often under-sampled (Chacoff et al., 2012; de Manincor, et al., 2020; Hegland et al., 2010; Nielsen & Bascompte, 2007; Olesen et al., 2011; Rivera-Hutinel et al., 2012; Sorensen et al., 2012; Stock et al., 2017; Vázquez et al., 2009). Which links are observed can relate to plant and pollinator generalism, morphological characteristics (Chacoff et al., 2012), or relative abundance (Blüthgen, 2010). While increasing sampling effort to sample all interactions might be prohibitively costly (Hegland et al., 2010), under-sampling potentially biases conclusions derived from these datasets, for example about levels of species specialization (Blüthgen, 2010; de Manincor et al., 2020; Petanidou et al., 2008; Rivera-Hutinel et al., 2012).

In this work, we explore whether realistic patterns of field under-sampling of plant-pollinator networks can be used to reconstruct full plant-pollinator networks when combined by cutting-edge link prediction methods (Van Kleunen et al., 2024). The majority of plant-pollinator network datasets have been sampled in the field by (a) recording a plant inventory at a site during a particular sampling period and then (b) recording observed visits of pollinators to plant species. Recent work has also constructed these networks via assessment of plants visited by insects that are captured in the field through clues obtained on insect bodies (e.g. DNA or pollen grains). Examination of pollen found on the bodies and specialized carrying structures of pollinators can be a more labor-intensive process than recording observed insect visits because it involves expert identification of pollen grains (for microscopic identifications) or comprehensive floral reference library (for DNA metabarcoding), but it generally provides a more complete view of the pollination network (Bosch et al., 2009; Cirtwill et al., 2024; Tourbez et al., 2023). Indeed, comparison of networks constructed via visit-based sampling and pollen-based sampling at the same sites have found that pollen-based sampling often identifies additional links and can also connect plants from a site’s plant inventory that were disconnected in the visit-based sampling network into the network, thus forming a more complete picture of pollination interactions at a site and revealing how visit-based sampling under-samples plant-pollinator links (de Manincor et al., 2020; Tourbez et al., 2023). Researchers have also noted that observed visits do not always indicate pollination interactions, and thus visit-based links may have additional biases (Ballantyne et al., 2015; Popic et al., 2013). Previous results comparing the networks produced from these two sampling strategies have had mixed results (Alarcón, 2010; Cirtwill et al., 2024; de Manincor et al., 2020; Tourbez et al., 2023), and it is still an open question whether networks sampled via visit observation alone can be used to study aspects of ecosystem stability relevant to decision making.

One emerging approach that could potentially help address under-sampling of interactions in visit-based sampling of plant-pollinator networks is using techniques for missing link prediction. Previous work has predicted missing links in plant-pollinator networks using pollinator and plant traits (e.g., Arroyo-Correa et al., 2021; Bartomeus, 2013; Bartomeus et al., 2016; Eklöf et al., 2013; Pichler et al., 2020) as well as aspects of network structure (e.g., Seo & Hutchinson, 2018; Stock et al., 2017; Terry & Lewis, 2020). We take advantage of advances in approaches for predicting missing links using stacked generalization, in which aspects of network structure (Ghasemian et al., 2020; Guimerà, 2020) and optionally node attributes representing species traits (Van Kleunen et al., 2024) are jointly considered in an ensemble for predicting missing links. We adapt this approach for use in bipartite networks with node attributes following expectations for link formation in plant-pollinator networks. We test the feasibility of this approach via application to a unique dataset in which plant-pollinator networks were sampled from three calcareous grasslands in France using both visit-based and pollen-based sampling strategies in the same month with high taxonomic precision (de Manincor et al., 2020).

We construct full networks for each site including all plants in the botanical inventory and all links identified via either sampling method (i.e., visits and pollen). To demonstrate the feasibility of using model-stacking for missing link prediction, we evaluate the ability of the models to re-predict uniform randomly removed links from these full networks, finding that the models are able to out-perform random baseline performance using either network structure or node attributes, which mirrors previous results for food webs (Van Kleunen et al., 2024). We next evaluate predicting missing links due to real-world patterns of link under-sampling represented by only including those links found with the visit-based sampling strategy. We find that predictive performance of the models is lower in this case, though still higher than random baselines across sites, indicating that one might feasibly partially reconstruct plausible full plant-pollinator networks from visit-based sampling. Thus, computational missing link prediction approaches have the potential to improve data quality for historical plant-pollinator networks, allowing for more robust analyses of pollination network stability over space and time. This is especially relevant as large databases of plant-pollinator networks and species traits are constructed (Aubouin et al., 2025; Lanuza et al., 2025).

An advantage of the stacked generalization approach is that it also allows one to investigate what aspects of network structure and what pollinator and plant traits drive better predictive performance for missing links, which can also provide insights into mechanisms driving link formation and observation (Ghasemian et al., 2020; Guimerà, 2020; Van Kleunen et al., 2024). Testing models using subsets of features, we found that across the three sites evaluated, structural predictors, which do not take into account node attributes, were the most important category driving predictive performance for the random under-sampling tests. Testing with real-world patterns of under-sampling, we found that node attributes became relatively more important for driving predictive performance across sites, in particular bee availability, plant availability, and bee pollen collection strategy, thus highlighting the utility of assembling trait data in cases of real-world sparse sampling of networks. Other specific categories of bee and plant traits (e.g., plant morphological traits) were more helpful for prediction at specific sites, thus also supporting the approach of learning to specialize a predictive model per network.

## Materials and Methods

### Study sites and plant-pollinator survey

We worked with plant-pollinator interaction data sampled from three calcareous grassland sites in France (Hauts-de-France, Regional natural reserve Riez de Noeux les Auxi, noted *R*; Normandie, Château Gaillard – le Bois Dumont, noted *CG*; Occitanie, Fourches, noted *F*; de Manincor et al., 2020). The three sites, of 1 ha each, are semi-natural protected areas included in the NATURA 2000 network. In each site, all flowering plants were identified at the species level in the field and their abundances recorded using the Braun-Blanquet coefficients of abundance-dominance (de Manincor et al., 2020b). For each flowering species we collected the anthers for pollen preparation in the laboratory. Pollinators, i.e. bees observed interacting with open flowers, were collected in the field and brought to the laboratory for preparation (pinning and pollen collection from bee’s bodies) and identification by expert taxonomists. The sampling took place in July 2016, the richest month of the year, during a one-day session of 4 hours in each site, for a total of 12 hours of observations (de Manincor et al., 2020). This study was part of a larger study (within the ANR ARSENIC project) where plant-pollinator interactions were recorded across 6 sites in France (2 sites per region, from the same three regions mentioned above) from April until October for two consecutive years. However, pollen preparation was only performed for the month of July 2016.

### Bee and plant trait data

We compiled trait data for the 69 unique bee species and 90 unique plant species across the 3 sites (further detail in Tables S1, S2).

Bee traits were retrieved from a database of European bee traits (established by ALARM, www.alarm-project.ufz.de, developed by STEP, www.STEP-project.net and maintained and updated by Stuart Roberts, Marshall et al., 2024), which combines bibliographic information and direct measurements made on specimens collected throughout Europe. For this study a subset of specimens from France was used. Information from this database was supplemented using the IUCN Red List of Threatened Species (https://www.iucnredlist.org/).

We classified bee traits into 4 groups:

1. **Foraging distance**: we used the ITD, which is the mean measurement (in mm) of the Inter Tegular Distance. The ITD can be used as a proxy for bee size related to the potential maximum foraging distance (Cane, 1987; Greenleaf et al., 2007).
2. **Pollen collection strategy**: we used information concerning the lecty (Cane, 2021; Cane & Sipes, 2006), tongue length (Cariveau et al., 2016; De Palma et al., 2015) and mode of pollen transport (Thorp, 2000).

a. *Lecty*: provides information on the bee diet, which can range from extremely specialized to extremely generalized. Bees can be identified as monolectic, i.e. foraging on a single plant species or a variety of plant species within a genus; oligolectic, i.e. foraging on pollen collected from a few species within one or a few families; polylectic, i.e. foraging on a variety of plant species and plant families.
b. *Tongue length*: this trait is given at the level of phylogenetic families, not at the species level. This trait distinguishes “long-tongue” from “short-tongue” bees. It informs on the bee’s ability to exploit resources from inflorescence types and can be used as a proxy for degree of floral specialization.
c. *Pollen transport*: represents the structural adaptation of bees to pollen transport, i.e. the way female bees collect and store pollen that will be used in the nest to feed the larvae. The pollen can be collected on their hind legs (including specialized structures such as the corbiculae for bumblebees and honeybees), on their body, or under the abdomen (typical for the Megachilidae family).
3. **Sociality**: indicates the species social behavior and can be used as a proxy for both reproductive strategy/success and foraging range (Grüter & Hayes, 2022; Wcislo & Fewell, 2017). In this study we only consider two categories, solitary or eusocial, because we excluded kleptoparasite and social parasite bees since they do not actively collect pollen. Solitary bees are characterized by individual female bees that construct the nest and provision the offspring alone, while eusocial bees are characterized by an overlap of generations, reproductive division of labor, and cooperative rearing of young.
4. **Bee Availability**: in this study we used bee relative abundance, i.e. number of sampled individuals for the same species, and phenology, i.e. presence of the bees during the season, to indicate fluctuations in the availability of bee populations to visit the flowers. For phenology, we used a proxy to indicate the presence of bees in relation to the month of our sampling (July): we created a binary indicator that is 0 if the species is in the first or last month of their season in July (also 0 if the bee phenology data indicates a 0 in July), and 1 if they are in the middle of their season as a proxy for the peak in relation to bee abundance potential.

Plant traits were retrieved from the database “Baseflor” (Julve, 1998) which provides information concerning floristic data and completed with information found on Tela Botanica (https://www.tela-botanica.org/), FloreAlpes (Le Driant, 2018) and literature (Tison et al., 2014; Tison & De Foucault, 2014). We classified plant traits in 3 groups:

1. **Reproduction strategy**: provides information on the mechanism governing pollination and mating (Barrett, 2010), and can be used as a proxy for pollination system specialization. We used four categories: apogamy, i.e. asexual reproduction, autogamy, i.e. self-fertilization, anemophily, i.e. wind pollination and entomophily, i.e. reproduction mediated by insects.
2. **Morphological traits** (associated with attractiveness): we identified morphological traits that may play a role in pollinator attractiveness and bee visitation patterns:

a. *Floral symmetry*: can be used by pollinators as a visual cue that provides information on the amount of resources (nectar or pollen, Giurfa et al., 1999; Ramírez, 2003). We distinguished actinomorphic, i.e. flowers with radial symmetry, and zygomorphic, i.e. flowers with bilateral symmetry.
b. *Floral display*: provides information on the inflorescence type from different floral classification systems (Ramírez, 2003). The floral display is directly associated with reproductive success. In this study we distinguished: solitary flowers, i.e. single inflorescence; capitulum or head, i.e. a dense vertically compressed inflorescence with sessile flowers on a receptacle, typical of the Asteraceae; umbel or corymb, i.e. a flat-topped or rounded inflorescence with the pedicels originating from a common point or a flat-topped raceme with elongate pedicels reaching the same level; spike or raceme, i.e. an elongate, unbranched, indeterminate inflorescence with either sessile or pedicelled flowers (categories from http://www.northernontarioflora.ca/); and other, a category encompassing all other types of inflorescences different from the previous.
c. *Floral color and color shade*: flower coloration is a visual signal used by plants to attract pollinators. In this study the floral color and color shade refer to the color perceived by humans and not the color perceived by bees, which is due to the wavelength-selective absorption by pigments and light scattering by structures inside the petals (Van Der Kooi et al., 2016, 2019). For flower color we distinguished blue, yellow, pink, green, and white flowers and light, neutral and dark for color shades.
3. **Flower availability**: similar to bee availability, in this study we used flower relative abundance and the phenology proxy to indicate the presence of flowering plants in relation to the month of our sampling (July, see above). For flower abundances, we used the abundances recorded during the botanic inventory and we transformed in flower cover percentages the classes of abundance-dominance of Braun-Blanquet (see de Manincor et al., 2020b for more details). For plant abundance, cover percentage was used as a proxy for the blooming potential (number of flowers produced) for each species.

### Versions of the plant-pollinator networks

The dataset is unique in that plant-pollinator networks for July 2016 were constructed for each site using two different methods for sampling plant-pollinator interactions. In the first method, plant-pollinator interactions were recorded via direct observation of bee visits to flowers during field sampling (the “Visit” set of interactions). In the second method, pollen was collected from the bodies of the same hand-net sampled wild bees and then identified by an expert in the laboratory to identify pollination interactions (the “Pollen” set of interactions). Interactions were only recorded for this study from non-parasitic female bees that actively collect pollen in the field. For the purposes of our analysis, we constructed a third version of the network that included the union of both of these sets of interactions for each site (hereafter referred to as the “Full” set of interactions), which we treated as the ground-truth plant-pollinator network for a site. For this analysis, we treated all interactions as binary. For the Full networks, we included as the set of nodes all bees for each site as well as all plants found in the plant inventory for a site, which meant that some plants were included that were fully disconnected from the Visit or Pollen networks. Although including disconnected plant nodes is atypical for constructing a plant-pollinator network for a site, we chose to do so due to our focus on predicting interactions from under-sampled networks in which it would be unknown *a priori* which plants would have pollination interactions with our pollinators of interest. We also excluded from the plant-pollinator networks plants that were found with the Pollen collection strategy which were not identified in the plant inventory, as these likely represented pollination interactions that occurred between the sampled bees and plants outside of the site, which cannot be identified as nodes *a priori* using the visit-based sampling method.

The number of nodes of each type, with disconnected plant nodes counted separately, as well as differences in the size of the interaction sets recorded with different strategies are shown in Table 1. Across the sites, the Pollen sampling method found more interactions for the set of bees, (31 more in the R site, 38 more in the CG site, and 107 more in the F site), supporting that it is generally a more comprehensive approach to sampling plant-pollinator interactions. However, it is of note that there were also interactions found with the Visit sampling method that were not found with the Pollen sampling method (9 more for R, 6 more for CG, and 22 more for F), indicating that this strategy either out-performed the Pollen strategy for detecting some interactions, or introduced superfluous interactions that did not represent true pollination interactions. These potentially superfluous interactions are included in our ground truth networks for our evaluation as their veracity could not be known from visit-based sampling alone. However, future work could attempt to identify these superfluous interactions.

**Table 1.** Plant-pollinator network properties. Number of nodes and edges in the plant-pollinator networks evaluated across the three sites and network versions. Counts of the differences between node and edge sets due to differences in sampling strategy are also indicated.

| Site | Network Version | Number of each node type |  |  | Total Nodes | Number Binary Edges |
| --- | --- | --- | --- | --- | --- | --- |
|  |  | *Bees* | *Connected plants* | *Disconnected plants* |  |  |
| Riez de Noeux les Auxi (R) | Visit | 19 | 12<br>(1 not in Pollen) |  | 31 | 42<br>(9 not in Pollen) |
|  | Pollen | 19 | 15<br>(4 not in Visit) |  | 34 | 64<br>(31 not in Visit) |
|  | Full | 19 | 16 | 15 | 50 | 73 |
| Château Gaillard – le Bois Dumont (CG) | Visit | 22 | 13 |  | 35 | 49<br>(6 not in Pollen) |
|  | Pollen | 22 | 18<br>(5 not in Visit) |  | 40 | 81<br>(38 not in Visit) |
|  | Full | 22 | 18 | 14 | 54 | 87 |
| Fourches (F) | Visit | 50 | 29<br>(4 plant nodes not in Pollen) |  | 79 | 98<br>(22 not in Pollen) |
|  | Pollen | 50 | 38<br>(13 not in Visit) |  | 88 | 183<br>(107 not in Visit) |
|  | Full | 50 | 42 | 10 | 102 | 205 |

### Stacking model approach for bipartite link prediction with node attributes

To predict missing links we used a stacked generalization approach adapted from Ghasemian et al., 2020, which was extended by Van Kleunen et al., 2024 to include node attributes. In missing link prediction, we assume that there is some true network G=(V,E) with a set of nodes V and a set of edges E but we only have access to an under-sampled network G’=(V,E’) where |E’|<|E|. Missing link predictions are reported as a ranking of all non-observed edges in G’ based on their predicted probability of being missing links. Thus, this is a binary classification task on all unobserved links to classify them as either “missing” (would be observed with increased sampling effort) or “forbidden” (cannot occur due to some temporal, spatial, or morphological constraint) (Olesen et al., 2011). In our analysis, the Full networks for each site were considered to be G. This approach uses a meta-learning model, as in Van Kleunen et al., 2024, to flexibly learn how to combine an ensemble of missing link prediction methods based on node attributes and network structure to predict missing links for a specific network (see Note S1 for more details on the training and test datasets used in the stacking model). We chose to use a random forest as our meta-learning model given its success in previous work across a broad corpus of network datasets (Ghasemian et al., 2020) and specifically in ecological networks (Van Kleunen et al., 2024), as well as previous work demonstrating the utility of random forests for link prediction in ecological networks (Llewelyn et al., 2023; Pichler et al., 2020; Van Kleunen et al., 2024; Wootton et al., 2024). We tailored this method to the bipartite context by adapting a suite of missing link prediction methods based on local network topology to the bipartite context (Note S1, Table S3), as well as adapting a suite of predictors based on node attributes for the bipartite context and based on trait-matching expectations for plant-pollinator networks (Note S1, Table S4). We trained and evaluated the model considering only possible bipartite links, i.e. links between the plant and pollinator levels, not within a single level (Note S1).

We performed two tests to evaluate missing link prediction performance. As a baseline, we first performed a test for each of the Full networks by repeatedly randomly removing 20% of the links in G to construct examples of G’. For each network, we repeated the process 5 times by randomly partitioning the set of links in G into 5 folds, removing each fold once to create G’ to ensure we performed tests removing all possible links. Link prediction performance was averaged across these 25 replications (5 repetitions × 5 folds) for each network. For our second test, we removed from the Full networks the set of interactions found with only the Pollen sampling strategy and not with the Visit sampling strategy, thus constructing G’ for each network via a real-world pattern of under-sampling. For this second test, results were averaged over 5 replications of missing link prediction to account for variations in performance introduced by random forest training and hyperparameter selection. To evaluate performance, we used two metrics: the area under the ROC (Receiver Operating Characteristics) curve (ROC-AUC) statistic (Ling et al., 2003) and the Precision-Recall AUC (PR-AUC) statistic (Poisot, 2023) calculated via the average precision score. In both cases, “better” models, insofar as they better predict the validation dataset, are the ones with higher AUC.

### Feature importance investigations

In order to evaluate the combination of plant and bee traits that leads to the best performance for missing link prediction, we repeated these two link prediction tests for each network while including different subsets of bee and plant traits. We tested 4 trait groups for bees (foraging distance, pollen collection strategy, sociality, and bee availability, 16 subsets including the empty set) and 3 trait groups for plants (reproduction strategy, morphological traits, and flower availability, 8 subsets including the empty set). We also tested predictive models with and without the set of topological predictors (called “structure”) derived from network structure alone. Overall, we tested 255 models for each site and test, representing all possible combinations of these groups.

Additionally, we investigated relative importance of individual predictors in these models by looking at feature importance scores. We looked at the Gini importance, which is derived from the training dataset during random forest training, and the permutation importance on the test dataset, which is calculated by repeatedly shuffling each feature and reporting the change in a performance metric, for which we use sensitivity and specificity.

## Results

### Top models for each site

We first report, for each site and task, the feature group subsets representing the top models ranked by ROC-AUC and PR-AUC performance (Table 2). ROC-AUC performances for the top models were higher with the random under-sampling for all sites, with a particularly large gap between under-sampling schemes for the R network (0.853 with random under-sampling and 0.661 with visit-based under-sampling). For PR-AUC, better performance was achieved predicting from the real-world patterns of under-sampling for the Fourches and Château Gaillard networks. We found that structure predictors were in all of the top models for the tests with random link removal. However, they were not for the tests with real-world Visit patterns of link removal. All of the groups of bee and plant traits showed up at least once in these top models, with variation across the sites and performance metrics.

**Table 2.** The groups of features included in the top performing models for each metric across the three sites for the two link prediction tests.

| Site | Under-sampling | ROC-AUC top model | ROC-AUC score | PR-AUC top model | PR-AUC score |
| --- | --- | --- | --- | --- | --- |
| Fourches | Random | Plant reproduction,<br>Plant morphological,<br>Bee pollen collection,<br>Structure | 0.806 | Plant reproduction,<br>Plant availability,<br>Bee pollen collection,<br>Bee availability,<br>Structure | 0.106 |
|  | Visit | Plant morphological,<br>Bee foraging distance,<br>Bee pollen collection,<br>Bee availability | 0.742 | Plant morphological,<br>Plant availability,<br>Bee ITD,<br>Bee pollen collection,<br>Bee availability | 0.133 |
| Riez de Noeux les Auxi | Random | Plant morphological,<br>Flower availability,<br>Bee pollen collection,<br>Structure | 0.853 | Plant reproduction,<br>Bee ITD,<br>Bee sociality,<br>Structure | 0.203 |
|  | Visit | Flower availability,<br>Bee pollen collection | 0.661 | Plant reproduction,<br>Bee ITD,<br>Bee pollen collection,<br>Bee sociality,<br>Bee availability | 0.136 |
| Château Gaillard – le Bois Dumont | Random | Bee pollen collection,<br>Bee sociality,<br>Bee availability,<br>Structure | 0.849 | Plant reproduction,<br>Bee ITD,<br>Bee pollen collection,<br>Structure | 0.165 |
|  | Visit | Plant reproduction,<br>Bee foraging distance,<br>Bee availability | 0.775 | Plant availability,<br>Bee ITD,<br>Bee pollen collection,<br>Bee sociality,<br>Bee availability | 0.198 |

### Comparing feature group predictive performance

In comparing the performance of models with and without groups of features, we saw for the test with random removal that the structure predictors drove a significant improvement in performance across all three sites for both performance metrics, as did bee availability except for the PR-AUC metric for the F site (Figure 1, Figure S1). Structure predictors significantly drove higher performance with real-world under-sampling for the PR-AUC metric across sites and the ROC-AUC metric for the F and CG sites but significantly decreased ROC-AUC performance for the R site, while bee availability significantly drove an increase in performance for all sites and both metrics. Model performance was notably improved more by structure predictors in random under-sampling than for real-world under-sampling. Bee pollen collection drove a significant improvement in performance for the F site for both metrics and for both random and real-world under-sampling, as well as for both other sites for the PR-AUC metric for real-world patterns of under-sampling and for the CG site for ROC-AUC performance under both sampling patterns. Plant availability also drove significant improvements for ROC-AUC performance across all three sites under random under-sampling, and for F and R sites under real-world under-sampling, as well as significant improvements in PR-AUC performance across all three sites for real-world under-sampling. Bee ITD drove significantly better performance for the CG network under random under-sampling for both metrics and for the PR-AUC metric for the R site under real-world under-sampling, but otherwise did not significantly impact performance. Plant morphological features drove better performance for the F network for real-world under-sampling for both metrics and for the ROC-AUC metric under random under-sampling, but otherwise did not improve performance and significantly decreased model performance for both other sites under real-world under-sampling. Bee sociality and plant reproduction strategy did not drive a significant improvement in performance across all sites, metrics, and sampling strategies, and bee sociality drove a significant decrease in performance for the PR-AUC metric for the R site.

**Figure 1.**
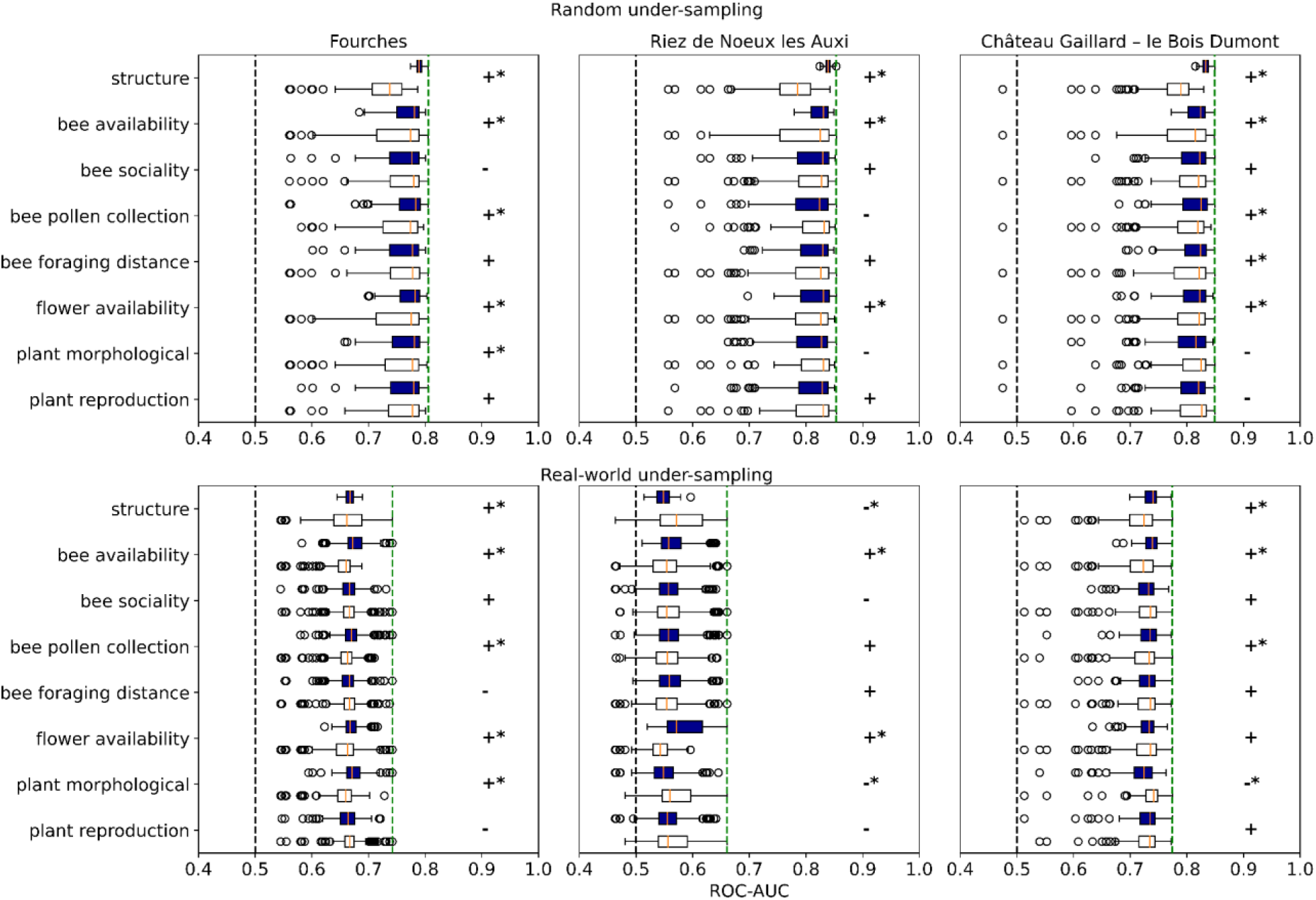
Comparison of model performance with and without groups of predictors (ROC-AUC). Results comparing average performance of models with (N=128, dark blue) and without (N=127, white) each group of predictors (structure, bee availability, bee sociality, bee pollen collection strategy, bee foraging distance, flower availability, plant morphological features, and plant reproduction strategy) across the three sites (Fourche, Riez de Noeux les Auxi, Château Gaillard – le Bois Dumont) based on the ROC-AUC metric for random 20% under-sampling and real-world patterns of under-sampling. Groups for which mean performance was better in models with that group of features rather than without are indicated with a + and otherwise with a -, and a significant difference in the mean performance between models with and without a group of features (p<0.05 via a two-sided t-test) are indicated with an asterisk. A black dashed line indicates the 0.5 baseline for ROC-AUC. A vertical green dashed line indicates the performance of the top model for that site and prediction task. PR-AUC performance is shown in Figure S1.

### Individual feature importances

Finally, we examined relative importance scores of individual features in the models with all predictors included to identify individual link prediction approaches that were relatively more useful in constructing ensemble predictions across sites (Figure 2, Figure S2, Figure S3). Across sites for both random and real-world under-sampling, some of the most influential predictors were the 9 structural predictors based on bipartite common neighbors (listed here with their abbreviations in Figure 2): bipartite common neighbors (BCN), bipartite resource allocation index (BRA), bipartite jaccard coefficient (BJC), bipartite Adamic/Adar index (BAA) (Daminelli et al., 2015), the Cannistraci variations of these same predictors (BCAR, BCRA, BCJC, BCAA) (Cannistraci et al., 2013) and bipartite community links (BLCL), indicating that missing link prediction based on local neighborhood structure was particularly helpful. The personalized page rank of plants for bees (PR-pers-bp in Figure 2) also consistently showed up amongst the most important predictors, as did the predictors based on low-rank approximations (LRA) of the bipartite adjacency matrix (bi-LRA, bi-LRA-dot in Figure 2). Plant and bee abundance stood out as particularly important amongst attribute-based predictors, thus suggesting that relative abundance drove the higher performance we observed for models containing bee and plant availability feature groups. Some of the derived trait-based predictors based on K-nearest neighbors (those prefixed with KNN in Figure 2) and Typical Neighbors (those prefixed with TN in Figure 2) were also amongst the most helpful attribute-based predictors. For example, KNN-b-D2 stood out as a relatively useful attribute-based feature for random under-sampling. This predictor was higher if the 3 nearest neighbor bees to the bee considered for a potential link (based on the Jaccard distances between the binary part of the bee attribute vectors) were neighbors with the plant considered for a potential link. Thus, these results support the use of such KNN and TN features, which combine information from bee and plant traits with local neighborhood structure, for missing link prediction in plant-pollinator networks.

**Figure 2.**
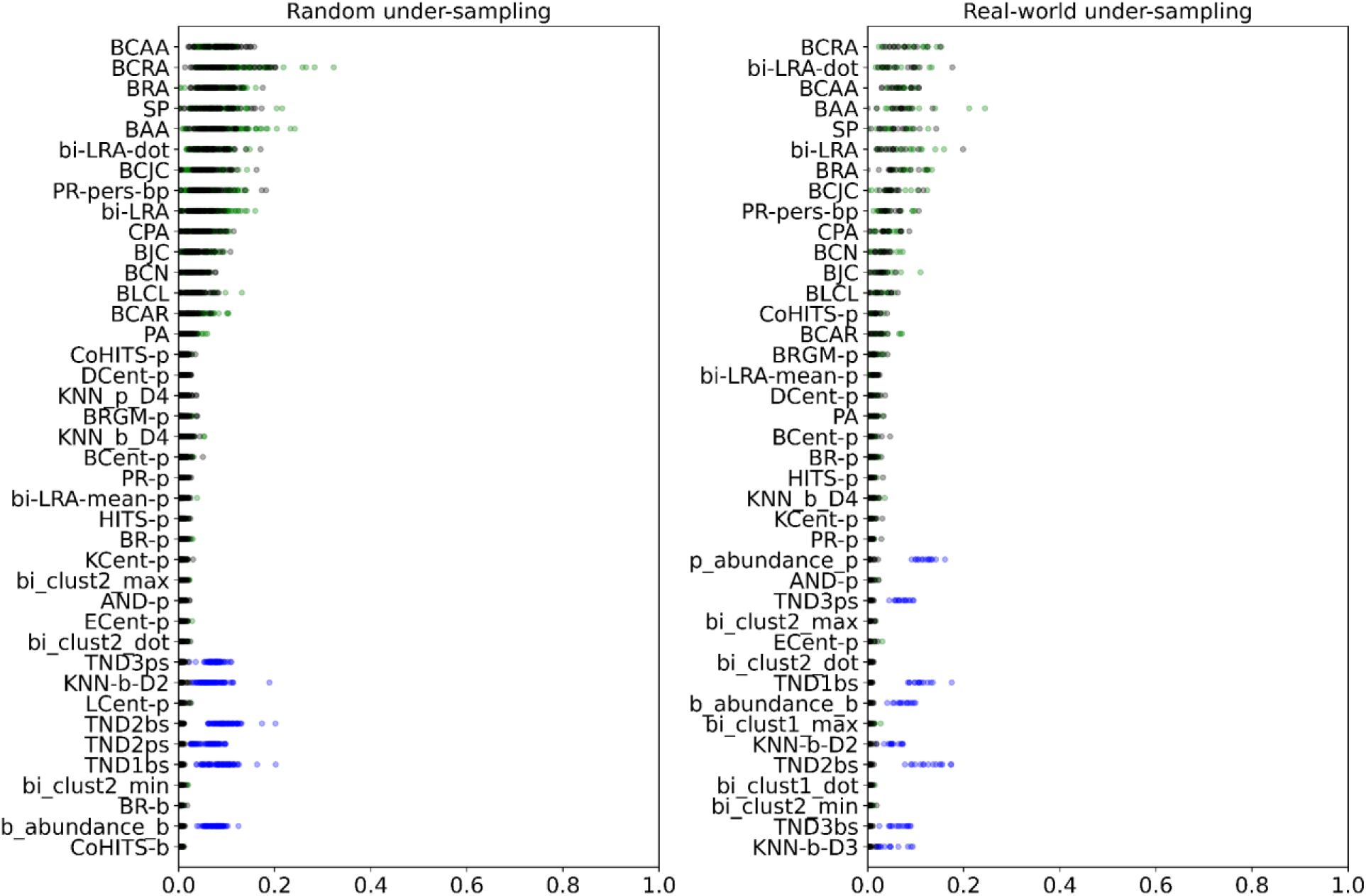
Gini importance results. The top 40 feature importance results are shown based on Gini importance (from the training dataset) in random and real-world under-sampling, summarized across all train/test splits across all sites for each feature for the model containing all features (in black) and ranked in descending order by mean importance. Results are contextualized by also plotting the relative importance of each feature in the models containing only structure predictors (in green) and the models containing no structure predictors (in blue). Comprehensive feature importance results are shown in Figure S2 (random under-sampling) and Figure S3 (real-world under-sampling). See Tables S1-S4 for feature definitions.

Binary predictors based on dummy variable encoding of categorical traits for bees and plants (e.g., pollen_transport_body_b, floral_color_dark_color_p) were generally ranked as having lower relative feature importance in the models. This result is likely partially attributable to the limitations of feature importance scores, where Gini importance scores are known to be biased towards higher feature importance attributed to numeric features (Nembrini et al., 2018) and permutation importance scores are known to have difficulty distinguishing important features that are highly correlated with other features in a model (Gregorutti et al., 2017). With these limitations in mind, the results for relative importance can still be compared between these binary traits. For example, flower color predictors seemed to have slightly higher relative Gini importance in the random under-sampling (Figure S2, Figure S3).

## Discussion

Our results on the imputation of missing links in plant-pollinator networks from three sites suggest that missing link prediction via a combination of network structure and traits is a potential strategy for re-constructing a more comprehensive version of pollination networks from the partially observed network constructed via direct observations of pollinator visits to plants. We found that structural predictors, with no additional species trait information, performed well at this task, indicating that this is also feasible in situations with limited or no trait data. However, collecting detailed plant and bee traits for use as node attributes improved predictive performance, and more so for predicting from real-world patterns of under-sampling than for predicting from random under-sampling. Visit-based sampling patterns could create structural biases (de Aguiar et al., 2019) – for example, previous analyses of this dataset indicated significant differences in species roles and specialization scores in visit-based networks compared to pollen-based networks (de Manincor et al., 2020). Structural predictors may improve predictive performance in some cases by effectively filling in information based on missing node attributes driving latent trait-based structure. However, at some level of down-sampling of links, learning these latent rules encoded in network structure might become impossible. Of note is the sparsity of this real-world under-sampling, which preserved the following percent of links in each network: 42/73 = 57.5% for R, 49/87 = 56.3% for CG, and 98/205 = 47.8% for F, whereas our tests with random under-sampling preserved 80% of the links. Prediction based on structural predictors should become less feasible with an increasing percentage of links removed, as less of the local neighborhood of nodes are preserved. Future work should test random under-sampling with increasing percentages of links removed to evaluate the degree to which this performance difference is driven by network sparsity.

In our model comparisons and investigation of relative feature importances, we saw overall that one could achieve good performance for predicting missing links using a combination of network structure and bee and plant availability. Our results also demonstrated differences between sites, supporting the stacking model approach, which effectively specializes a predictive model by site. For example, for the Fourches site, located in Occitanie in southern France and the most diverse site for both bees and plants, plant morphological features were useful in predicting missing links, whereas this was not the case for the other two sites. This was a site in which there were many specialized bee species, and so plant traits might have driven interaction predictions more for this site than the other two sites. The stacking model approach is also flexible in that it could use different traits as node attributes if traits other than those we evaluated are available for a site.

The way in which we processed data for our analysis made some assumptions. In adding bee and plant traits as attributes, we collected trait values from general sources and we assumed that these would be representative for species at our sites. This might not always be the case - for example, plant phenology can shift per site based on factors such as soil quality, precipitation, climate, pollination activity, or competition (Cleland et al., 2006; Dunne et al., 2003; Pashalidou et al., 2020; Peñuelas et al., 2004; Wolf et al., 2017). We made an assumption that the union of the Visit and Pollen links in our dataset could be used as the ground-truth plant-pollinator network for a site in evaluating model performance. However, this data might itself include missing links, and the visit-based networks can also include superfluous links not present in the pollen networks (Tourbez et al., 2023). Further, we made a choice to exclude all plants found via Pollen-based sampling that were external to the site based on the prediction task we were interested in in which these plant nodes would not be known. However, these are open systems and these external plant nodes might still play a role in ecosystem stability. We also focused only on bees as they are the key pollinators in these systems, but other pollinators may also play some role in stability. More generally, it is important to keep in mind that the set of “missing interactions” heavily depends on some artificial closure of an open system. This closure is at once temporal (“are links missing if the interaction occurred a week or a month ago?”), spatial (“do interactions with plants in nearby fields count?”), and taxonomic (“are links with shrubs or trees missing or simply overlooked?”).

In this study we only looked at predicting binary interactions. However, visit-based networks are often constructed as weighted networks, in which the number of visits observed is taken as a link strength between a plant and pollinator. While this information was included indirectly in our models via bee relative abundance, future work could attempt to predict link weights. Because bees move, it is more difficult to estimate bee relative abundance than plant relative abundance at a site. We ultimately chose to use bee relative abundance in our models, though we note that the bee relative abundance value was not independent of interaction sampling in the same way as plant relative abundance, as this value was recorded based on the number of bees sampled at a site when sampling for interactions (i.e., one sampled bee led to one recorded interaction in the original, weighted network version of the dataset). Thus, while a plant might have high relative abundance and no or low interactions in the dataset, a bee with high relative abundance would necessarily have had many interactions sampled at a site (although it’s possible that these produced a lower number of binary interactions if multiple individuals of the same bee species were observed interacting with the same plant species).

In real-world under-sampling we expect traits to relate to missingness of links in two ways: (a) by constraining which interactions are possible (trait-matching) and (b) by influencing which links are more likely to be missing (i.e., non-random under-sampling based on node attributes). For instance, plant and pollinator phenologies constrain whether species can interact – if plants do not flower at the same time the insect species is actively collecting pollen, there is no way they can interact; on the other hand, interactions between small pollinators and any plant are more likely be missing, especially when both species are rare, since more conspicuous pollinators can be more easily captured. While we considered these components together when predicting from real-world patterns of under-sampling, future work could attempt to distinguish these two components of missingness. Further, traits encoded as node attributes could also be related to erroneously recorded links, if for example a taxonomic group is more challenging to identify correctly, which we do not yet address in our approach. When choosing which node attributes to include in the models we considered both aspects of missingness. For example, whether plant relative abundances can act as a species trait constraining interactions is an open question. The relationship between species relative abundance and interaction patterns is challenging to untangle, as it has been noted that more abundant species often appear more generalist and the causal direction of this relationship is unknown (Fort et al., 2016). However, there are cases in which pollen specialists for a specific plant will always visit only that plant even when it is rare. For example, this was observed with *Rophites* bees in the Fourches site, which only visited rare *Stachys* flowers. While we were unable to untangle these effects, we included plant abundances in our models because we thought that plant relative abundances would at least affect whether or not interactions were observed by the research team (de Manincor et al., 2020b). Future work could investigate how the nodes participating in the empirical missing links in our dataset (those found in the Pollen network but not the Visit) as well as those participating in the empirical links that were likely superfluous (those found in the Visit network but not Pollen) differed in their trait distributions from those of the connected species at a site overall.

Moreover, we do not explicitly yet tie our results related to binary plant-pollinator networks to implications for ecological stability. It is likely that binary interactions alone do not contain all of the information necessary for assessing how pollinator declines will affect biodiversity and ecosystem services. For example, some interactions might involve more efficient pollination than others, that is a plant’s true set of pollinators from the perspective of ecological stability might be a subset of its observed interaction partners (Sargent & Otto, 2006). Ultimately we are interested in to what extent plant-pollinator networks sampled via visit observation can be used to assess ecological stability. While distinguishing more consequential pollination interactions might be difficult without additional ecological data, future work could investigate whether the prediction techniques such as those we have demonstrated in this work could aid in gaining a more comprehensive understanding of plant-pollinator network stability.

## Acknowledgements

Financial support for data collection was provided by the ANR projects ARSENIC (grant no. 14-CE02-0012) and NGB (grant no. 17-CE32-0011). Lucy Van Kleunen acknowledges funding from the Chateaubriand Fellowship. We also thank Stéphane Robin, Julien Chiquet, Sophie Donnet, and other members of the ANR EcoNet working group (ANR-18-CE02-0010) for the fruitful discussions, and Julian Resasco, Leana Zoller, Daniel B. Larremore, Elizabeth Bradley and Amir Ghasemian for discussions of the in progress work.

## References

1. Alarcón, R. (2010). Congruence between visitation and pollen-transport networks in a California plant–pollinator community. Oikos, 119(1), 35–44. 10.1111/j.1600-0706.2009.17694.x

2. Arroyo-Correa, B., Bartomeus, I., & Jordano, P. (2021). Individual-based plant–pollinator networks are structured by phenotypic and microsite plant traits. Journal of Ecology, 109(8), 2832–2844. 10.1111/1365-2745.13694

3. Aubouin, L., Genoud, D., Givord-Coupeau, B., Tercerie, S., Gargominy, O., Geslin, B., & Schatz, B. (2025). BeeFunc, a comprehensive trait database for French bees. Scientific Data, 12(1), 1302. 10.1038/s41597-025-05626-0

4. Ballantyne, G., Baldock, K. C. R., & Willmer, P. G. (2015). Constructing more informative plant–pollinator networks: Visitation and pollen deposition networks in a heathland plant community. Proceedings of the Royal Society B: Biological Sciences, 282(1814), 20151130. 10.1098/rspb.2015.1130

5. Barrett, S. C. H. (2010). Understanding plant reproductive diversity. Philosophical Transactions of the Royal Society B: Biological Sciences, 365(1537), 99–109. 10.1098/rstb.2009.0199

6. Bartomeus, I. (2013). Understanding Linkage Rules in Plant-Pollinator Networks by Using Hierarchical Models That Incorporate Pollinator Detectability and Plant Traits. PLoS ONE, 8(7), e69200. 10.1371/journal.pone.0069200

7. Bartomeus, I., Gravel, D., Tylianakis, J. M., Aizen, M. A., Dickie, I. A., & Bernard-Verdier, M. (2016). A common framework for identifying linkage rules across different types of interactions. Funct. Ecol., 30(12), 1894–1903. 10.1111/1365-2435.12666

8. Blüthgen, N. (2010). Why network analysis is often disconnected from community ecology: A critique and an ecologist’s guide. Basic and Applied Ecology, 11(3), 185–195. 10.1016/j.baae.2010.01.001

9. Bosch, J., Martín González, A. M., Rodrigo, A., & Navarro, D. (2009). Plant-pollinator networks: Adding the pollinator’s perspective. Ecology Letters, 12(5), 409–419. 10.1111/j.1461-0248.2009.01296.x

10. Cane, J. H. (1987). Estimation of Bee Size Using Intertegular Span (Apoidea). Journal of the Kansas Entomological Society, 60(1), 145–147.

11. Cane, J. H. (2021). A brief review of monolecty in bees and benefits of a broadened definition. Apidologie, 52(1), 17–22. 10.1007/s13592-020-00785-y

12. Cane, J. H., & Sipes, S. (2006). Characterizing floral specialization by bees: Analytical methods and a revised lexicon for oligolecty. In Waser NM, Ollerton J (eds) Plant-pollinator interactions: From specialization to generalization. (1st ed., pp. 99–122). The University of Chicago Press.

13. Cannistraci, C. V., Alanis-Lobato, G., & Ravasi, T. (2013). From link-prediction in brain connectomes and protein interactomes to the local-community-paradigm in complex networks. Scientific Reports, 3(1), 1613. 10.1038/srep01613

14. Cariveau, D. P., Nayak, G. K., Bartomeus, I., Zientek, J., Ascher, J. S., Gibbs, J., & Winfree, R. (2016). The Allometry of Bee Proboscis Length and Its Uses in Ecology. PLOS ONE, 11(3), e0151482. 10.1371/journal.pone.0151482

15. Chacoff, N. P., Vázquez, D. P., Lomáscolo, S. B., Stevani, E. L., Dorado, J., & Padrón, B. (2012). Evaluating sampling completeness in a desert plant-pollinator network: Sampling a plant-pollinator network. Journal of Animal Ecology, 81(1), 190–200. 10.1111/j.1365-2656.2011.01883.x

16. Cirtwill, A. R., Wirta, H., Kaartinen, R., Ballantyne, G., Stone, G. N., Cunnold, H., Tiusanen, M., & Roslin, T. (2024). Flower-visitor and pollen-load data provide complementary insight into species and individual network roles. Oikos, 2024(4), e10301. 10.1111/oik.10301

17. Cleland, E. E., Chiariello, N. R., Loarie, S. R., Mooney, H. A., & Field, C. B. (2006). Diverse responses of phenology to global changes in a grassland ecosystem. Proceedings of the National Academy of Sciences, 103(37), 13740–13744. 10.1073/pnas.0600815103

18. Daminelli, S., Thomas, J. M., Durán, C., & Vittorio Cannistraci, C. (2015). Common neighbours and the local-community-paradigm for topological link prediction in bipartite networks. New Journal of Physics, 17(11), 113037. 10.1088/1367-2630/17/11/113037

19. de Aguiar, M. A. M., Newman, E. A., Pires, M. M., Yeakel, J. D., Boettiger, C., Burkle, L. A., Gravel, D., Guimarães, P. R., O’Donnell, J. L., Poisot, T., Fortin, M.-J., & Hembry, D. H. (2019). Revealing biases in the sampling of ecological interaction networks. PeerJ, 7, e7566. 10.7717/peerj.7566

20. de Manincor, N., Hautekèete, N., Mazoyer, C., Moreau, P., Piquot, Y., Schatz, B., Schmitt, E., Zélazny, M., & Massol, F. (2020). How biased is our perception of plant-pollinator networks? A comparison of visit- and pollen-based representations of the same networks. Acta Oecologica, 105, 103551. 10.1016/j.actao.2020.103551

21. de Manincor, N., Hautekeete, N., Piquot, Y., Schatz, B., Vanappelghem, C., & Massol, F. (2020b). Does phenology explain plant–pollinator interactions at different latitudes? An assessment of its explanatory power in plant–hoverfly networks in French calcareous grasslands. Oikos, 129(5), 753–765. 10.1111/oik.07259

22. De Palma, A., Kuhlmann, M., Roberts, S. P. M., Potts, S. G., Börger, L., Hudson, L. N., Lysenko, I., Newbold, T., & Purvis, A. (2015). Ecological traits affect the sensitivity of bees to land-use pressures in E uropean agricultural landscapes. Journal of Applied Ecology, 52(6), 1567–1577. 10.1111/1365-2664.12524

23. Dunne, J. A., Harte, J., & Taylor, K. J. (2003). SUBALPINE MEADOW FLOWERING PHENOLOGY RESPONSES TO CLIMATE CHANGE: INTEGRATING EXPERIMENTAL AND GRADIENT METHODS. Ecological Monographs, 73(1), 69–86. 10.1890/0012-9615(2003)073[0069:SMFPRT]2.0.CO;2

24. Eklöf, A., Jacob, U., Kopp, J., Bosch, J., Castro-Urgal, R., Chacoff, N. P., Dalsgaard, B., de Sassi, C., Galetti, M., Guimarães, P. R., Lomáscolo, S. B., Martín González, A. M., Pizo, M. A., Rader, R., Rodrigo, A., Tylianakis, J. M., Vázquez, D. P., & Allesina, S. (2013). The dimensionality of ecological networks. Ecology Letters, 16(5), 577–583. 10.1111/ele.12081

25. Fontaine, C., Dajoz, I., Meriguet, J., & Loreau, M. (2005). Functional Diversity of Plant–Pollinator Interaction Webs Enhances the Persistence of Plant Communities. PLoS Biology, 4(1), e1. 10.1371/journal.pbio.0040001

26. Fort, H., Vázquez, D. P., & Lan, B. L. (2016). Abundance and generalisation in mutualistic networks: Solving the chicken-and-egg dilemma. Ecology Letters, 19(1), 4–11. 10.1111/ele.12535

27. Ghasemian, A., Hosseinmardi, H., Galstyan, A., Airoldi, E. M., & Clauset, A. (2020). Stacking models for nearly optimal link prediction in complex networks. Proceedings of the National Academy of Sciences, 117(38), 23393–23400. 10.1073/pnas.1914950117

28. Gill, R. J., Baldock, K. C. R., Brown, M. J. F., Cresswell, J. E., Dicks, L. V., Fountain, M. T., Garratt, M. P. D., Gough, L. A., Heard, M. S., Holland, J. M., Ollerton, J., Stone, G. N., Tang, C. Q., Vanbergen, A. J., Vogler, A. P., Woodward, G., Arce, A. N., Boatman, N. D., Brand-Hardy, R., … Potts, S. G. (2016). Protecting an Ecosystem Service: Approaches to Understanding and Mitigating Threats to Wild Insect Pollinators. In Advances in Ecological Research (Vol. 54, pp. 135–206). Elsevier. 10.1016/bs.aecr.2015.10.007

29. Giurfa, M., Dafni, A., & Neal, P. R. (1999). Floral Symmetry and Its Role in Plant-Pollinator Systems. International Journal of Plant Sciences, 160(S6), S41–S50. 10.1086/314214

30. Greenleaf, S. S., Williams, N. M., Winfree, R., & Kremen, C. (2007). Bee foraging ranges and their relationship to body size. Oecologia, 153(3), 589–596. 10.1007/s00442-007-0752-9

31. Gregorutti, B., Michel, B., & Saint-Pierre, P. (2017). Correlation and variable importance in random forests. Statistics and Computing, 27(3), 659–678. 10.1007/s11222-016-9646-1

32. Grüter, C., & Hayes, L. (2022). Sociality is a key driver of foraging ranges in bees. Current Biology, 32(24), 5390–5397.e3. 10.1016/j.cub.2022.10.064

33. Guimerà, R. (2020). One model to rule them all in network science? Proceedings of the National Academy of Sciences, 117(41), 25195–25197. 10.1073/pnas.2017807117

34. Hegland, S. J., Dunne, J., Nielsen, A., & Memmott, J. (2010). How to monitor ecological communities cost-efficiently: The example of plant–pollinator networks. Biological Conservation, 143(9), 2092–2101. 10.1016/j.biocon.2010.05.018

35. Julve, P. (1998). Baseflore. Index botanique, écologique et chorologique de la flore de France. [Dataset]. 10.13140/RG.2.1.3561.5601

36. Keyes, A. A., McLaughlin, J. P., Barner, A. K., & Dee, L. E. (2021). An ecological network approach to predict ecosystem service vulnerability to species losses. Nature Communications, 12(1), 1586. 10.1038/s41467-021-21824-x

37. Lanuza, J. B., Knight, T. M., Montes-Perez, N., Glenny, W., Acuña, P., Albrecht, M., Artamendi, M., Badenhausser, I., Bennett, J. M., Biella, P., Bommarco, R., Cappellari, A., Castro, S., Clough, Y., Colom, P., Costa, J., Cyrille, N., De Manincor, N., Dominguez-Lapido, P., … Bartomeus, I. (2025). EUPPOLLNET: A European Database of Plant-Pollinator Networks. Global Ecology and Biogeography, 34(2), e70000. 10.1111/geb.70000

38. Le Driant, F. (2018). *Flore Alpes* [Dataset]. www.florealpes.com/

39. Ling, C. X., Huang, J., & Zhang, H. (2003). AUC: A Better Measure than Accuracy in Comparing Learning Algorithms. In Y. Xiang & B. Chaib-draa (Eds.), Advances in Artificial Intelligence (Vol. 2671, pp. 329–341). Springer Berlin Heidelberg. 10.1007/3-540-44886-1_25

40. Llewelyn, J., Strona, G., Dickman, C. R., Greenville, A. C., Wardle, G. M., Lee, M. S. Y., Doherty, S., Shabani, F., Saltré, F., & Bradshaw, C. J. A. (2023). Predicting predator–prey interactions in terrestrial endotherms using random forest. Ecography, e06619. 10.1111/ecog.06619

41. Marshall, L., Leclercq, N., Carvalheiro, L. G., Dathe, H. H., Jacobi, B., Kuhlmann, M., Potts, S. G., Rasmont, P., Roberts, S. P. M., & Vereecken, N. J. (2024). Understanding and addressing shortfalls in European wild bee data. Biological Conservation, 290, 110455. 10.1016/j.biocon.2024.110455

42. Nembrini, S., König, I. R., & Wright, M. N. (2018). The revival of the Gini importance? Bioinformatics, 34(21), 3711–3718. 10.1093/bioinformatics/bty373

43. Nielsen, A., & Bascompte, J. (2007). Ecological networks, nestedness and sampling effort. Journal of Ecology, 95(5), 1134–1141. 10.1111/j.1365-2745.2007.01271.x

44. Okuyama, T., & Holland, J. N. (2008). Network structural properties mediate the stability of mutualistic communities. Ecology Letters, 11(3), 208–216. 10.1111/j.1461-0248.2007.01137.x

45. Olesen, J. M., Bascompte, J., Dupont, Y. L., Elberling, H., Rasmussen, C., & Jordano, P. (2011). Missing and forbidden links in mutualistic networks. Proceedings of the Royal Society B: Biological Sciences, 278(1706), 725–732. 10.1098/rspb.2010.1371

46. Ollerton, J., Winfree, R., & Tarrant, S. (2011). How many flowering plants are pollinated by animals? Oikos, 120(3), 321–326. 10.1111/j.1600-0706.2010.18644.x

47. Pashalidou, F. G., Lambert, H., Peybernes, T., Mescher, M. C., & De Moraes, C. M. (2020). Bumble bees damage plant leaves and accelerate flower production when pollen is scarce. Science, 368(6493), 881–884. 10.1126/science.aay0496

48. Peñuelas, J., Filella, I., Zhang, X., Llorens, L., Ogaya, R., Lloret, F., Comas, P., Estiarte, M., & Terradas, J. (2004). Complex spatiotemporal phenological shifts as a response to rainfall changes. New Phytologist, 161(3), 837–846. 10.1111/j.1469-8137.2004.01003.x

49. Petanidou, T., Kallimanis, A. S., Tzanopoulos, J., Sgardelis, S. P., & Pantis, J. D. (2008). Long-term observation of a pollination network: Fluctuation in species and interactions, relative invariance of network structure and implications for estimates of specialization: High plasticity in plant-pollinator networks. Ecology Letters, 11(6), 564–575. 10.1111/j.1461-0248.2008.01170.x

50. Pichler, M., Boreux, V., Klein, A., Schleuning, M., & Hartig, F. (2020). Machine learning algorithms to infer trait-matching and predict species interactions in ecological networks. Methods in Ecology and Evolution, 11(2), 281–293. 10.1111/2041-210X.13329

51. Poisot, T. (2023). Guidelines for the prediction of species interactions through binary classification. Methods in Ecology and Evolution, 14(5), 1333–1345. 10.1111/2041-210X.14071

52. Popic, T. J., Wardle, G. M., & Davila, Y. C. (2013). Flower-visitor networks only partially predict the function of pollen transport by bees. Austral Ecology, 38(1), 76–86. 10.1111/j.1442-9993.2012.02377.x

53. Potts, S. G. (with Imperatriz-Fonseca, V. L., Ngo, H. T., Biesmeijer, J. C., Breeze, T. D., Dicks, L. V., Garibaldi, L. A., Hill, R., Settele, J., & Vanbergen, A. J.). (2016). The assessment report on pollinators, pollination and food production: Summary for policymakers. Secretariat of the Intergovernmental Science-Policy Platform on Biodiversity and Ecosystem Services.

54. Ramírez, N. (2003). Floral specialization and pollination: A quantitative analysis and comparison of the Leppik and the Faegri and van der Pijl classification systems. TAXON, 52(4), 687–700. 10.2307/4135542

55. Rivera-Hutinel, A., Bustamante, R. O., Marín, V. H., & Medel, R. (2012). Effects of sampling completeness on the structure of plant–pollinator networks. Ecology, 93(7), 1593–1603. 10.1890/11-1803.1

56. Ross, S. R. P.-J., Arnoldi, J.-F., Loreau, M., White, C. D., Stout, J. C., Jackson, A. L., & Donohue, I. (2021). Universal scaling of robustness of ecosystem services to species loss. Nature Communications, 12(1), 5167. 10.1038/s41467-021-25507-5

57. Sánchez-Bayo, F., & Wyckhuys, K. A. G. (2019). Worldwide decline of the entomofauna: A review of its drivers. Biological Conservation, 232, 8–27. 10.1016/j.biocon.2019.01.020

58. Sargent, R. D., & Otto, S. P. (2006). The Role of Local Species Abundance in the Evolution of Pollinator Attraction in Flowering Plants. The American Naturalist, 167(1).

59. Seo, E., & Hutchinson, R. A. (2018). Predicting links in plant-pollinator interaction networks using latent factor models with implicit feedback. Thirty-Second AAAI Conference on Artificial Intelligence, 808–815.

60. Sorensen, P. B., Damgaard, C. F., Strandberg, B., Dupont, Y. L., Pedersen, M. B., Carvalheiro, L. G., Biesmeijer, J. C., Olsen, J. M., Hagen, M., & Potts, S. G. (2012). A method for under-sampled ecological network data analysis: Plant-pollination as case study. Journal of Pollination Ecology, 6. 10.26786/1920-7603(2011)18

61. Stock, M., Poisot, T., Waegeman, W., & De Baets, B. (2017). Linear filtering reveals false negatives in species interaction data. Scientific Reports, 7(1), 45908. 10.1038/srep45908

62. Terry, J. C. D., & Lewis, O. T. (2020). Finding missing links in interaction networks. Ecology, 101(7). 10.1002/ecy.3047

63. Thébault, E., & Fontaine, C. (2010). Stability of Ecological Communities and the Architecture of Mutualistic and Trophic Networks. Science, 329(5993), 853–856. 10.1126/science.1188321

64. Thorp, R. W. (2000). The collection of pollen by bees. 222, 211–223.

65. Tison, J. M., & De Foucault, B. (2014). Flora gallica: Flore de France.

66. Tison, J. M., Jauzein, P., & Michaud, H. (2014). Flore de la France méditerranéenne continentale (Vol. 1). Naturalia Publications.

67. Tourbez, C., Gómez-Martínez, C., González-Estévez, M. Á., & Lázaro, A. (2023). Pollen analysis reveals the effects of uncovered interactions, pollen-carrying structures, and pollinator sex on the structure of wild bee–plant networks. Insect Science, 1744–7917.13267. 10.1111/1744-7917.13267

68. Traveset, A., & Richardson, D. M. (2014). Mutualistic Interactions and Biological Invasions. Annual Review of Ecology, Evolution, and Systematics, 45(1), 89–113. 10.1146/annurev-ecolsys-120213-091857

69. Valdovinos, F. S. (2022, August 18). Effects of network structure and adaptive foraging on the ecosystem services provided by plant-pollinator systems. ESACSEE 2022, Montreal.

70. Valdovinos, F. S., Berlow, E. L., Moisset de Espanés, P., Ramos-Jiliberto, R., Vázquez, D. P., & Martinez, N. D. (2018). Species traits and network structure predict the success and impacts of pollinator invasions. Nature Communications, 9(1), 2153. 10.1038/s41467-018-04593-y

71. Van Der Kooi, C. J., Dyer, A. G., Kevan, P. G., & Lunau, K. (2019). Functional significance of the optical properties of flowers for visual signalling. Annals of Botany, 123(2), 263–276. 10.1093/aob/mcy119

72. Van Der Kooi, C. J., Elzenga, J. T. M., Staal, M., & Stavenga, D. G. (2016). How to colour a flower: On the optical principles of flower coloration. Proceedings of the Royal Society B: Biological Sciences, 283(1830), 20160429. 10.1098/rspb.2016.0429

73. Van Kleunen, L., Dee, L. E., Wootton, K. L., Massol, F., & Clauset, A. (2024). Predicting missing links in food webs using stacked models and species traits. Ecology. 10.1101/2024.11.22.624890

74. Vanbergen, A. J., & Initiative, T. I. P. (2013). Threats to an ecosystem service: Pressures on pollinators. Frontiers in Ecology and the Environment, 11(5), 251–259. 10.1890/120126

75. Vázquez, D. P., Blüthgen, N., Cagnolo, L., & Chacoff, N. P. (2009). Uniting pattern and process in plant–animal mutualistic networks: A review. Annals of Botany, 103(9), 1445–1457. 10.1093/aob/mcp057

76. Wcislo, W., & Fewell, J. H. (2017). Sociality in Bees. In Comparative Social Evolution (1st ed., pp. 50–83). Cambridge University Press. 10.1017/9781107338319.004

77. Wolf, A. A., Zavaleta, E. S., & Selmants, P. C. (2017). Flowering phenology shifts in response to biodiversity loss. Proceedings of the National Academy of Sciences, 114(13), 3463–3468. 10.1073/pnas.1608357114

78. Wootton, K. L., Guillaume Blanchet, F., Liston, A., Nyman, T., Riggi, L. G. A., Kopelke, J., Roslin, T., & Gravel, D. (2024). Layer-specific imprints of traits within a plant–herbivore–predator network – complementary insights from complementary methods. Ecography, 2024(4), e07028. 10.1111/ecog.07028

