## Supplementary Information for "Understanding and addressing under-sampling in plant-pollinator networks using stacked models for missing link prediction"

**Table of Contents**

Supplemental Figures

Figure S1. Comparison of model performance with and without groups of predictors (PR-AUC).

Figure S2. Relative importance of individual predictors for predicting missing links (random under-sampling).

Figure S3. Relative importance of individual predictors for predicting missing links (real-world under-sampling).

Supplemental Methods

Note 1. Stacking model used for missing link prediction.

Supplemental Tables

Table S1. Trait information included as node attributes for each bee species.

Table S2. Trait information included as node attributes for each plant species.

Table S3. Structure-based predictors included in the stacking model.

Table S4. Attribute-based predictors included in the stacking model.

Supplemental Figures

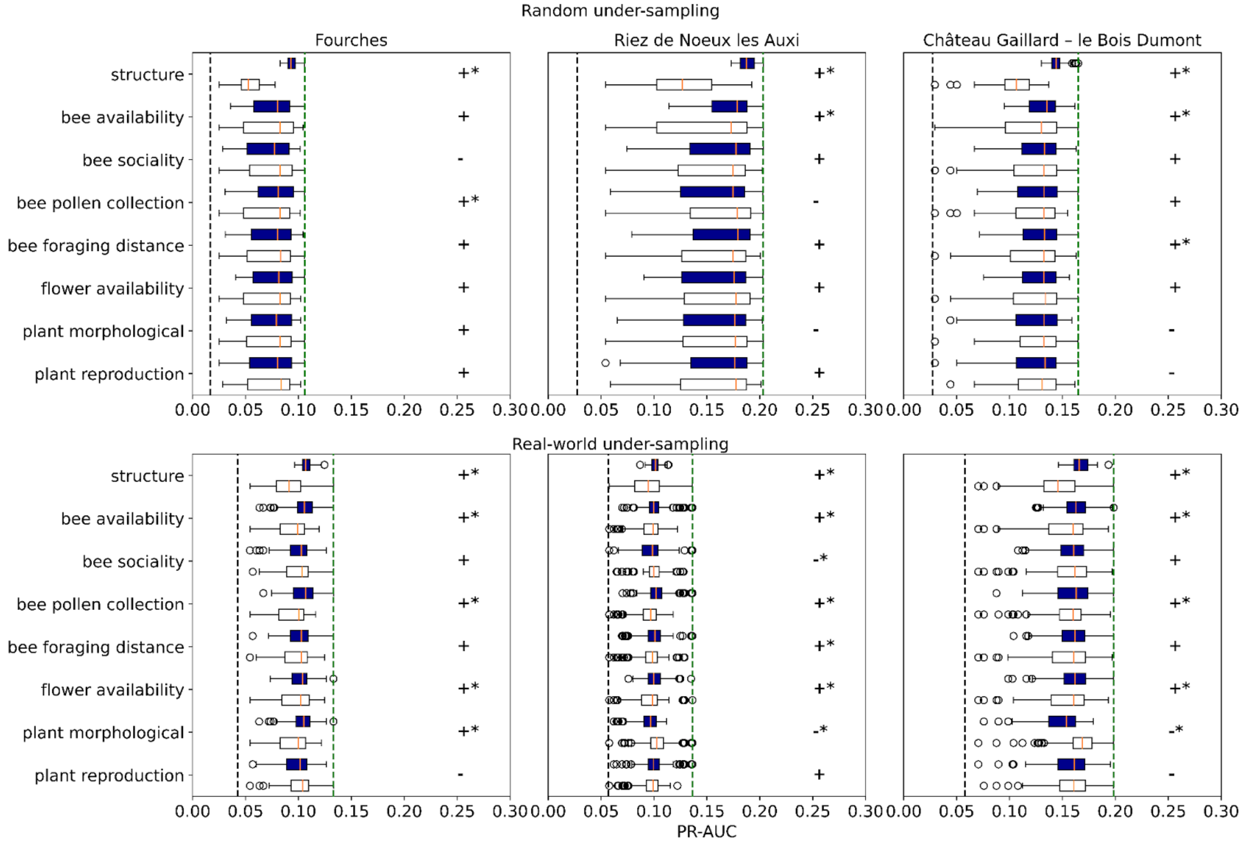


**Figure S1. Comparison of model performance with and without groups of predictors (PR-AUC).** Results comparing average performance of models with (N=128, dark blue) and without (N=127, white) each group of predictors (structure, bee availability, bee sociality, bee pollen collection strategy, bee foraging distance, flower availability, plant morphological features, and plant reproduction strategy) across the three sites (Fourche, Riez de Noeux les Auxi, Château Gaillard – le Bois Dumont) based on the PR-AUC metric for random 20% under-sampling and real-world patterns of under-sampling. Groups for which mean performance was better in models with that group of features rather than without are indicated with a + and otherwise with a -, and a significant difference in the mean performance between models with and without a group of features (p<0.05 via a two-sided t-test) are indicated with an asterisk. A black dashed line indicates the baseline for the PR-AUC metric (this value is dataset-specific based on the proportion of the positive class). A vertical green dashed line indicates the performance of the top model for that site and prediction task.

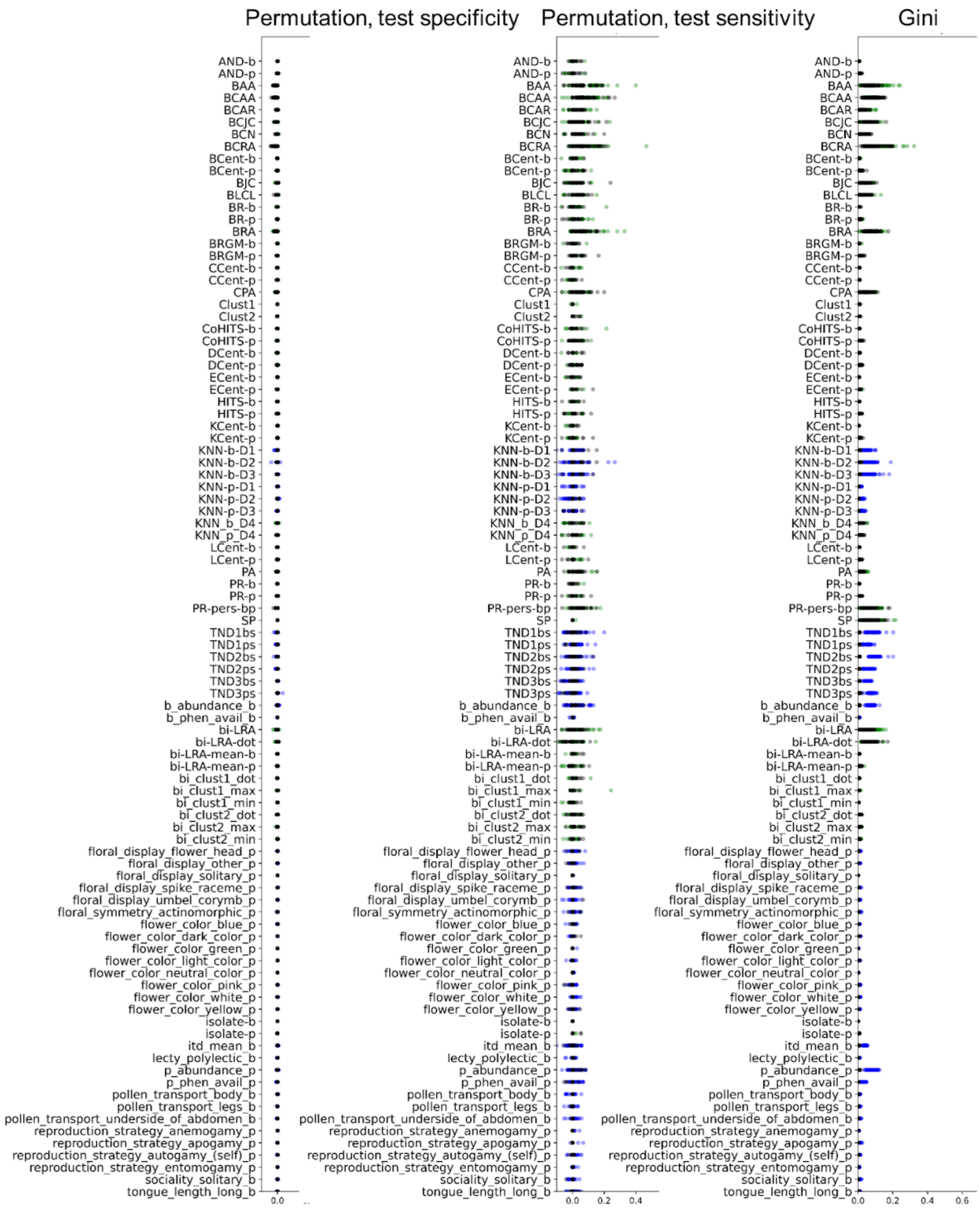


**Figure S2. Relative importance of individual predictors for predicting missing links (random under-sampling)**. Feature importance results are shown based on Gini importance (from the training dataset) and permutation importance (for sensitivity and specificity on the test dataset) for random under-sampling, summarized across all train/test splits across all sites for each feature for the model containing all features (in black). Results are contextualized by also plotting the relative importance of each feature in the models containing only structure predictors (in green) and the models containing no structure predictors (in blue).

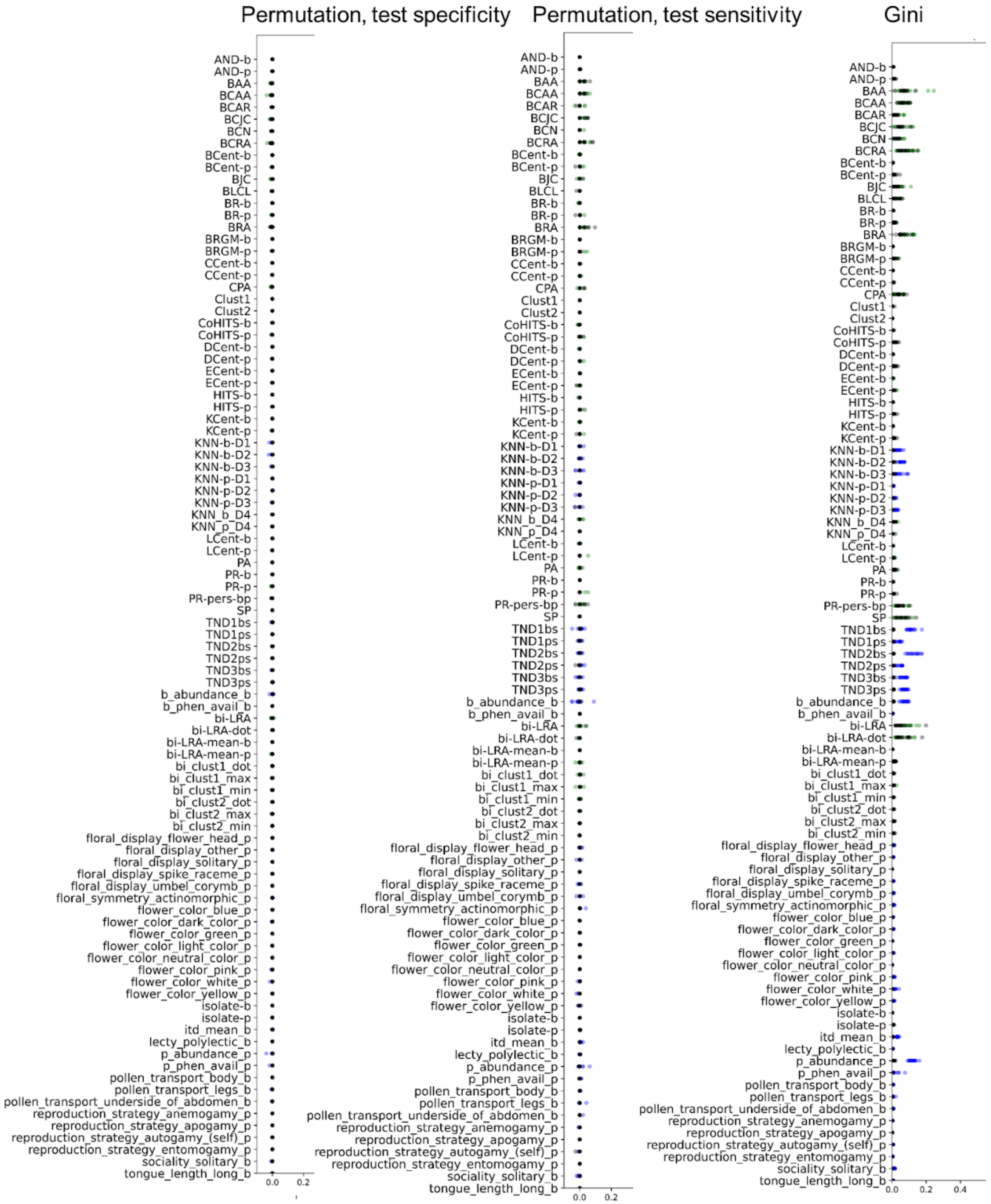


**Figure S3. Relative importance of individual predictors for predicting missing links (real-world under-sampling)**. Feature importance results are shown based on Gini importance (from the training dataset) and permutation importance (for sensitivity and specificity on the test dataset) for real-world under-sampling, summarized across all train/test splits across all sites for each feature for the model containing all features (in black). Results are contextualized by also plotting the relative importance of each feature in the models containing only structure predictors (in green) and the models containing no structure predictors (in blue).

Supplemental Methods

**Note 1. Developing a stacking model used for bipartite missing link prediction for plant-pollinator networks.**

For missing link prediction, we used a method based on stacked generalization (Ghasemian et al., 2020). Previous results on ecological networks have shown that this approach can be used to predict missing links by learning how to combine node attributes, such as species traits, and local topological properties of pairs of nodes encoded in structural predictors (Van Kleunen et al., 2024).

In a bipartite network G=(V, E) with node set V and edge set E, the node set V can be split into two distinct sets, A and B, where links can only form between nodes of these two different types, not between nodes of the same type. For the missing link prediction problem, we assume that we only have access to a subset of E, E' in which E \ E' are the missing links. To train a model to distinguish missing from true non-links, E' is further down-sampled to E'', with E' \ E'' forming the positive class of examples of missing links and all non-links in G'=(V, E') taken as negative examples of missing links. To decrease the imbalance between the positive and negative classes for small networks, in this work we also add as examples to the positive class all links in E'', which encode trait-matching signals. The size of the training dataset is thus equal to the number of non-links in G'' = |A|*|*B*|-|*E''*| *plus* |*E''*|*, or* |*A*|*|B|. For each node pair *b* (bee node) and node *p* (plant node), predictors are calculated from G'' to form features for this training dataset, based on their attribute vectors and their interactions in G''. A meta-level model is trained on this dataset and then applied to a test dataset formed from all of the non-links in G', of size |A|*|B|-|E'|, with predictors derived from G', to rank these by their probability of being missing links. In our case we followed prior work (Ghasemian et al., 2020; Van Kleunen et al.,

2024) in using a random forest model as our meta-level model, with hyperparameters selected via 5-fold cross validation optimizing for PR-AUC following advice in Poisot, 2023.

Predictors based on node attributes are listed in Table S4. All node attributes were included as raw values. Pairwise attribute-based predictors used in previous work (Van Kleunen et al., 2024) based on the distance between node attribute vectors were inappropriate in this context in which node pairs involve nodes of two different types and thus do not share the same attributes. We replaced these predictors with what we call "typical neighbor" (*TN*) predictors based on the distance between the typical neighbor vector of node *b* and the attribute vector of node *p* (*TND{n}bs*) and the distance between the attribute vector of node *b* and the typical neighbor vector of node *p* (*TND{n}ps*). The typical neighbor vector of a node was constructed by aggregating the attribute vectors of all of a node's neighbors by taking the mode of each position for binary values (with ties broken randomly), and means for numeric values. For disconnected nodes with no neighbors, the typical neighbor was constructed based on the aggregation of all nodes of the other node type. The intuition behind these predictors is that node *b* and node *p* are more likely to interact if they are respectively similar to the existing interaction partners of these nodes. Most of the attributes we considered for bees (except ITD Mean and abundance) and all of the attributes we considered for plants except abundance were categorical attributes that had been transformed into binary values. We considered two distance metrics between the binary parts of the attribute vectors, *D1*: the Manhattan distance and *D2*: the Jaccard distance, leading to 4 predictors (*TND1bs*, *TND1ps*, *TND2bs*, *TND2ps*). When numeric attributes were included for either node type we added an additional distance metric *D3* which was the Euclidean distance between the numeric part of the attribute vectors (min-max normalized), which is simply the absolute value of the difference in the case of only one numeric attribute (*TND3bs*, *TND3ps*). For example, when ITD Mean was included as the only numeric attribute for bees, this distance became the difference between node *p*'s typical neighbor ITD Mean value and node *b*'s ITD Mean value.

We also added 6 K-nearest neighbor (*KNN*) (Desjardins-Proulx et al., 2017) predictors based on node attributes (*KNN-b-D1, KNN-b-D2, KNN-p-D1, KNN-p-D2, KNN-b-D3, KNN-p-D3*). With a similar motivation as the typical neighbor predictors, *KNN* predictors are constructed by finding the *K* nearest neighbors of node *b* (or node *p*) and then recording what fraction of node *b*'s (or node *p*'s) *K* nearest neighbors contain node *p* (or node *b*) in their neighbor sets (Van Kleunen et al., 2024). We used the same 3 distance metrics *D1*, *D2*, and *D3* to determine *K* nearest neighbors and set *K* to 3. *KNN-b-D3* and *KNN-p-D3* were only included if bees or plants had numeric attributes included respectively. Note that both *TN* and *KNN* attribute-based predictors include information on the neighbor sets of nodes and thus are not based solely on node attributes but also include auxiliary information about local network structure.

We tested each model including an additional 53 predictors based entirely on network structure (Ghasemian et al., 2020) (Table S3). Previous results found best performance for link prediction in ecological networks by combining node attribute predictors with network structure predictors (Van Kleunen et al., 2024). All of the predictors we considered were topological predictors, which can easily be easily calculated for a node pair in a network based on the network adjacency matrix and do not require complex model parameter tuning. The set of topological predictors were chosen for inclusion in the models based on their ability to predict missing links as an ensemble in prior work and then adjusted for the bipartite context as appropriate. We also considered the biological interpretability of these predictors as they related to mechanisms that might drive plant-pollinator link formation. Many of these topological predictors are highly correlated and as a whole they provide a comprehensive picture of the local neighborhood of a node pair representing a potential missing link.

Amongst the predictors we included two preferential attachment predictors (*PA, CPA*) (Daminelli et al., 2015), based on the assumption that generalist bees and generalist pollinators are more likely to interact with each other. We additionally added 2 binary predictors indicating specifically if nodes were completely disconnected from the food web (e.g., were of degree 0; *isolate-b, isolate-p*), which might occur with link down-sampling during model training for small networks and was particularly relevant in our application due to the difference in plant node sets between Visit and Pollen sampling strategies. These predictors are based on the assumption that fully disconnected nodes are less likely to have missing links. We also included 2 predictors based on average neighbor degrees (*AND-b, AND-p*) (Barrat et al., 2004), which extend the assumption about degree determining links to the neighbors of nodes.

We included 6 node centrality metrics based on one-mode projections of the bipartite network (*ECent-b, ECent-p, KCent-b, KCent-p, LCent-b, LCent-p*) (Goh et al., 2001; Hagberg et al., 2008; Katz, 1953; Newman, 2001). This strategy separately calculates centrality scores for each node type in the bipartite network, though the strategy of one-mode projection can lose some of the information from the entire bipartite network (Yang et al., 2020). We thus also included 6 centrality metrics calculated on the entire bipartite network with calculations adjusted for bipartite degree expectations (*CCent-b*, *CCent-p*, *DCent-b, DCent-p, BCent-b, BCent-p*) (Borgatti & Halgin, 2014). Centrality predictors could encode scenarios in which nodes that are more or less central players in the network are more likely to have missing links and also be used to qualify other predictor values.

Similarly we followed prior work in including ranking predictors, including a PageRank value calculated on one-mode projections of the bipartite network (*PR-b*, *PR-p*) and a personalized page rank value calculated on the full network (*PR-pers-bp*) (Page et al., 1999) as well as 8 ranking predictors specifically developed for ranking on bipartite networks (*BR-b, BR-p, HITS-b, HITS-p, CoHITS-b, CoHITS-p, BRGM-b, BRGM-p*) (Yang et al., 2020). Similar to centrality predictors, ranking predictors can help with prediction if nodes that are higher in a latent hierarchy within the network are more likely to be missing links as well as be used to qualify the values of other predictors.

4 predictors were included based on a low-rank approximation (LRA) via singular value decomposition of the bipartite adjacency matrix (*bi-LRA*, *bi-LRA-dot*, *bi-LRA-mean-b*, *bi-LRA-mean-p*). The low rank approximation essentially represents each node with a low dimensional vector in a shared latent trait space based on node interactions. The *bi-LRA* and *bi-LRA-do*t predictors are based on the proximity of the latent trait vectors of the two nodes within this space, while the *bi-LRA-mean-b* and *bi-LRA-mean-p* predictors are similar to the typical neighbor predictors but in latent trait space (and vice versa) (Cukierski et al., 2011; Dalla Riva & Stouffer, 2016; Koren et al., 2009; Massol et al., 2021). We chose the rank to use in the LRA using the heuristic from Zhu & Ghodsi, 2006.

Successful link prediction has often been achieved based on the assumption that there are clusters or tightly connected local neighborhoods within a bipartite network. 1 predictor included based on these assumptions was the shortest paths (*SP*) predictor, which indicates the proximity of a node pair in a network. We also included 9 predictors based on the concept of bipartite common neighbors from Daminelli et al., 2015 (B*CN, BLCL, BRA, BAA, BCRA, BCAA, BJC, BCAR, BCJC*), which all generally assume that nodes are more likely to interact if they have many links in the same local neighborhood of the network. Finally, we added 2 KNN predictors (*KNN-b-D4, KNN-p-D4*) based only on network structure, in which *D4* was the Jaccard distance between the neighbor sets of the two nodes, thus assuming that a bee would be more likely to interact with a plant (and vice versa) if many bees it shared plant neighbors with also did. Nodes with empty neighbor sets, for which the Jaccard similarity is not defined, were considered at a distance of 0 given these nodes would be considered similar in our application (e.g., two disconnected plant nodes). We also included 6 predictors based on three versions of the bipartite specific clustering coefficient, which encodes to what extent each node is in a densely clustered section of the network, which might also indicate it is more likely to have missing links (*bi_clust1_min, bi_clust2_min, bi_clust1_max, bi_clust2_max, bi_clust1_dot, bi_clust2_dot*) (Latapy et al., 2008). Finally, we added 2 predictors (*Clust1, Clust2*) using spectral clustering. Spectral clustering attempts to identify tightly connected groups in a network based on KMeans clustering of the normalized eigenvectors of the Laplacian of the network adjacency matrix (Chung, 1997; Ng et al., 2002; Von Luxburg, 2007; White & Smyth, 2005). The connected part of the network was clustered with disconnected nodes excluded and treated together as a separate group. We chose the number of clusters using the spectral gap heuristic based on the Laplacian of the network adjacency matrix (Von Luxburg, 2007; White & Smyth, 2005), choosing the top two options for the number of clusters to construct *Clust1* and *Clust2* respectively. The predictor values were considered 1 if nodes *b* and *p* were in the same identified cluster, based on the assumption that nodes in the same tightly-connected cluster would be more likely to interact, and 0 otherwise.

Supplemental Tables

**Table S1. Trait information included as node attributes for each bee species.**

| **Trait name** | **Possible values** | **Trait group** |
|---|---|---|
| ITD Mean | Float (mm) | Foraging Distance |
| Lecty | Binary indicator:<br>*polylectic = 1;*<br>*oligolectic = 0* | Pollen collection strategy |
| Pollen transport | Binary indicator per category (3, can be multiple):<br>*legs;*<br>*body;*<br>*underside of abdomen* | Pollen collection strategy |
| Tongue Length | Binary indicator:<br>*long = 1;*<br>*short = 0* | Pollen collection strategy |
| Sociality | Binary indicator:<br>*solitary = 1;*<br>*eusocial = 0* | Sociality |
| Phenology | Binary indicator:<br>*Bee is in the middle of its season = 1;*<br>*Bee is in the first or last month or outside of its season = 0* | Bee Availability |
| Relative Abundance | Integer (number of individuals) | Bee Availability |

**Table S2. Trait information included as node attributes for each plant species.**

| Trait name | Possible values | Trait group |
|---|---|---|
| Reproduction Strategy | Binary indicator per category (4):<br>*apogamy;*<br>*autogamy (self);*<br>*anemogamy;*<br>*entomogamy* | Reproduction strategy |
| Floral Display | Binary indicator per category (5):<br>*solitary;*<br>*flower head;*<br>*umbel corymb;*<br>*spike raceme;*<br>*other* | Plant morphology |
| Flower Color | Binary indicator per category (5):<br>*blue;*<br>*yellow;*<br>*pink;*<br>*green;*<br>*white* | Plant morphology |
| Flower Color Shade | Binary indicator per category (3):<br>*light color;*<br>*neutral color;*<br>*dark color* | Plant morphology |
| Floral Symmetry | Binary indicator:<br>*actinomorphic = 1;*<br>*zygomorphic = 0* | Plant morphology |
| Phenology | Binary indicator:<br>*Plant is in the middle of its season = 1;*<br>*Plant is in the first or last month of its season = 0* | Flower availability |
| Relative Abundance | Float (percent cover) | Flower availability |

**Table S3. Structure predictors included in the stacking model.** Predictors were implemented using helper functions from the NetworkX software library (Hagberg et al., 2008).

| Resolution | Abbreviation | Definition | References |
|---|---|---|---|
| Pairwise | PA | Preferential attachment (degree product) between *b* and *p* | (Daminelli et al., 2015) |
| Pairwise | CPA | Cannistraci variation of preferential attachment | (Cannistraci et al., 2013; Daminelli et al., 2015) |
| Node-level | isolate-b, isolate-p | Whether *b* or *p* are isolates (fully disconnected) | |
| Node-level | AND-b, AND-p | Average neighbor degrees for *b* and *p* | (Barrat et al., 2004) |
| Node-level | ECent-b, ECent-p | Eigenvector centralities for *b* and *p*, calculated separately on one-mode projections of the graph | See NetworkX documentation for a review (Hagberg et al., 2008). |
| Node-level | KCent-b, KCent-p | Katz centralities for *b* and *p*, calculated separately on one-mode projections of the graph | (Katz, 1953) |

| | | | |
|---|---|---|---|
| Node-level | LCent-b,<br>LCent-p | Load centralities for *b* and *p*, calculated separately on one-mode projections of the graph | (Goh et al., 2001; Newman, 2001) |
| Node-level | CCent-b,<br>CCent-p | Bipartite node closeness centrality for nodes *b* and *p* | (Borgatti & Halgin, 2014) |
| Node-level | DCent-b,<br>DCent-p | Bipartite node degree centrality for nodes *b* and *p* | (Borgatti & Halgin, 2014) |
| Node-level | BCent-b,<br>BCent-p | Bipartite node betweenness centrality for nodes *b* and *p* | (Borgatti & Halgin, 2014) |
| Pairwise | PR-pers-bp | *p*th entry of the personalized page rank of node *b* | (Page et al., 1999) |
| Node-level | PR-b,<br>PR-p | Page rank values for *b* and *p* | (Page et al., 1999) |
| Node-level | BR-b,<br>BR-p | BiRank values for *b* and *p* | (Yang et al., 2020) |
| Node-level | HITS-b,<br>HITS-p | HITS rankings for *b* and *p* | (Yang et al., 2020) |
| Node-level | CoHITS-b,<br>CoHITS-p | CoHITS rankings for *b* and *p* | (Yang et al., 2020) |

| | | | |
|---|---|---|---|
| Node-level | BRGM-b, BRGM-p | BGRM rankings for *b* and *p* | (Yang et al., 2020) |
| Pairwise | bi-LRA | Entry *b*, *p* in low rank approximation (LRA) of the bipartite network adjacency matrix via singular value decomposition (SVD). | (Cukierski et al., 2011; Dalla Riva & Stouffer, 2016), Rank choice (Zhu & Ghodsi, 2006). |
| Pairwise | bi-LRA-dot | Where SVD creates three matrices, L', S, and R'; the dot product of the *b*th row of L' and the *p*th row of R'. | (Cukierski et al., 2011; Dalla Riva & Stouffer, 2016) adjusted for bipartite, Rank choice (Zhu & Ghodsi, 2006). |
| Pairwise | bi-LRA-mean-b | The mean value of LRA[*b*, *j*] for all *b*'s neighbors *j*. | (Cukierski et al., 2011), adjusted for bipartite. |
| Pairwise | bi-LRA-mean-p | The mean value of LRA[*i*, *p*] for all *p*'s neighbors *i*. | (Cukierski et al., 2011), adjusted for bipartite. |
| Pairwise | SP | Shortest path between *b*, *p* | (Liben-Nowell & Kleinberg, 2007) |

| | | | |
|---|---|---|---|
| Pairwise | BCN | Number of bipartite common neighbors | (Daminelli et al., 2015) |
| Pairwise | BLCL | Number of bipartite local community links | (Daminelli et al., 2015) |
| Pairwise | BRA | Bipartite resource allocation index | (Daminelli et al., 2015) |
| Pairwise | BAA | Bipartite Adamic Adar index | (Daminelli et al., 2015) |
| Pairwise | BCRA | Cannistraci variation of bipartite resource allocation index | (Cannistraci et al., 2013; Daminelli et al., 2015) |
| Pairwise | BCAA | Cannistraci variation of bipartite Adamic Adar index | (Cannistraci et al., 2013; Daminelli et al., 2015) |

| | | | |
|---|---|---|---|
| Pairwise | BJC | Bipartite Jaccard coefficient | (Daminelli et al., 2015) |
| Pairwise | BCAR | Cannistraci variation of bipartite common neighbors | (Cannistraci et al., 2013; Daminelli et al., 2015) |
| Pairwise | BCJC | Cannistraci variation of bipartite Jaccard coefficient | (Cannistraci et al., 2013; Daminelli et al., 2015) |
| Pairwise | KNN-b-D4 | The fraction of node *b*'s K nearest neighbors of the same type (where D4 is the Jaccard similarity based on *p*-neighbor sets) of which node *p* is a neighbor | (Van Kleunen et al., 2024) |
| Pairwise | KNN-p–D4 | The fraction of node *p*'s K nearest neighbors of the same type (where D4 is the Jaccard similarity based on *b*-neighbor sets) of which node *b* is a neighbor | (Van Kleunen et al., 2024) |

| | | | |
|---|---|---|---|
| Node-level | bi_clust1_min,<br>bi_clust2_min | Bipartite clustering coefficient for nodes *b* (1) and *p* (2), using mode min | (Latapy et al., 2008) |
| Node-level | bi_clust1_max,<br>bi_clust2_max | Bipartite clustering coefficient for nodes *b* (1) and *p* (2), using mode max | (Latapy et al., 2008) |
| Node-level | bi_clust1_dot,<br>bi_clust2_dot | Bipartite clustering coefficient for nodes *b* (1) and *p* (2), using mode dot | (Latapy et al., 2008) |
| Pairwise | Clust1 | Binary indicator - whether *b* and *p* were found in the same cluster found via spectral clustering (for the top number of clusters). | (Von Luxburg, 2007; White & Smyth, 2005);<br>Code: (Chiquet, 2024), accessed 2024. |
| Pairwise | Clust2 | Binary indicator - whether *b* and *p* were found in the same cluster found via spectral clustering (for the second best number of clusters). | (Von Luxburg, 2007; White & Smyth, 2005);<br>Code: (Chiquet, 2024), accessed 2024. |

**Table S4. Attribute-based predictors included in the stacking model.** Predictors were implemented using helper functions from the NetworkX software library (Hagberg et al., 2008). KNN predictors are based on Van Kleunen et al., 2024.

| **Resolution** | **Abbreviation** | **Definition** |
|---|---|---|
| Node-level | {x}_b,{y}_p | Raw attribute values (repeated for all bee attributes *x* and all plant attributes *y*. |
| Pairwise | TND1bs | The Manhattan distance between the binary attribute vectors of the typical neighbor of node *b* and node *p* |
| Pairwise | TND1ps | he Manhattan distance between the binary part of the attribute vectors of the typical neighbor of node *p* and node *b* |
| Pairwise | TND2bs | The Jaccard distance between the binary attribute vectors of the typical neighbor of node *b* and node *p* |
| Pairwise | TND2ps | he Jaccard distance between the binary part of the attribute vectors of the typical neighbor of node *p* and node *b* |
| Pairwise | TND3bs | The Euclidean distance between the numeric parts of the attribute vectors of the typical neighbor of node *b* and node *p* (only included if numeric attributes were included for plants) |

| | | |
|---|---|---|
| Pairwise | TND3ps | The Euclidean distance between the min-max normalized numeric parts of the attribute vectors of the typical neighbor of node *p* and node *b* (only included if numeric attributes were included for bees) |
| Pairwise | KNN-b-D1 | The fraction of node *b*'s *K* nearest neighbors of the same type (as determined by the Manhattan distance between the binary part of the attribute vectors) of which node *p* is a neighbor |
| Pairwise | KNN-p-D1 | The fraction of node *p*'s *K* nearest neighbors of the same type (as determined by the Manhattan distance between the binary attribute vectors) of which node *b* is a neighbor |
| Pairwise | KNN-b-D2 | The fraction of node *b*'s *K* nearest neighbors of the same type (as determined by the Jaccard distance between the binary part of the attribute vectors) of which node *p* is a neighbor |
| Pairwise | KNN-p-D2 | The fraction of node *p*'s *K* nearest neighbors of the same type (as determined by the Jaccard distance between the binary attribute vectors) of which node *b* is a neighbor |
| Pairwise | KNN-b-D3 | The fraction of node *b*'s *K* nearest neighbors of the same type (as determined by the Euclidean distance between the min-max normalized numeric part of the attribute vectors) of which node *p* is a neighbor. |

| | | |
|---|---|---|
| Pairwise | KNN-p-D3 | The fraction of node $p$'s $K$ nearest neighbors of the same type (as determined by the Euclidean distance between the min-max normalized numeric part of the attribute vectors) of which node $b$ is a neighbor. |

### References


Barrat, A., Barthélemy, M., Pastor-Satorras, R., & Vespignani, A. (2004). The architecture of complex weighted networks. *Proceedings of the National Academy of Sciences*, *101*(11), 3747–3752. https://doi.org/10.1073/pnas.0400087101

Borgatti, S. P., & Halgin, D. S. (2014). Analyzing Affiliation Networks. In J. Scott & P. Carrington, *The SAGE Handbook of Social Network Analysis* (pp. 417–433). SAGE Publications Ltd. https://doi.org/10.4135/9781446294413.n28

Cannistraci, C. V., Alanis-Lobato, G., & Ravasi, T. (2013). From link-prediction in brain connectomes and protein interactomes to the local-community-paradigm in complex networks. *Scientific Reports*, *3*(1), 1613. https://doi.org/10.1038/srep01613

Chiquet, J. (2024, Accessed). *Hierarchical and Spectral methods for Graph clustering*. https://jchiquet.github.io/MAP566/docs/mixture-models/map566-lecture-graph-clustering-part1.html

Chung, F. R. (1997). *Spectral graph theory* (Vol. 92). American Mathematical Soc.

Cukierski, W., Hamner, B., & Yang, B. (2011). Graph-based features for supervised link prediction. *The 2011 International Joint Conference on Neural Networks*, 1237–1244. https://doi.org/10.1109/IJCNN.2011.6033365

Dalla Riva, G. V., & Stouffer, D. B. (2016). Exploring the evolutionary signature of food webs' backbones using functional traits. *Oikos*, *125*(4), 446–456. https://doi.org/10.1111/oik.02305

Daminelli, S., Thomas, J. M., Durán, C., & Vittorio Cannistraci, C. (2015). Common neighbours and the local-community-paradigm for topological link prediction in bipartite networks. *New Journal of Physics*, *17*(11), 113037. https://doi.org/10.1088/1367-2630/17/11/113037

Desjardins-Proulx, P., Laigle, I., Poisot, T., & Gravel, D. (2017). Ecological interactions and the Netflix problem. *PeerJ*, *5*, e3644. https://doi.org/10.7717/peerj.3644

Ghasemian, A., Hosseinmardi, H., Galstyan, A., Airoldi, E. M., & Clauset, A. (2020). Stacking models for nearly optimal link prediction in complex networks. *Proceedings of the National Academy of Sciences*, *117*(38), 23393–23400. https://doi.org/10.1073/pnas.1914950117

Goh, K.-I., Kahng, B., & Kim, D. (2001). Universal Behavior of Load Distribution in Scale-Free Networks. *Physical Review Letters*, *87*(27), 278701. https://doi.org/10.1103/PhysRevLett.87.278701

Hagberg, A., Swart, P., & Chult, D. (2008). *Exploring network structure, dynamics, and function using NetworkX.* (No. No. LA-UR-08-05495; LA-UR-08-5495.). Los Alamos National Lab (LANL).

Katz, L. (1953). A new status index derived from sociometric analysis. *Psychometrika*, *18*(1), 39–43. https://doi.org/10.1007/BF02289026

Koren, Y., Bell, R., & Volinsky, C. (2009). Matrix Factorization Techniques for Recommender Systems. *Computer*, *42*(8), 30–37. https://doi.org/10.1109/MC.2009.263

Latapy, M., Magnien, C., & Vecchio, N. D. (2008). Basic notions for the analysis of large two-mode networks. *Social Networks*, *30*(1), 31–48. https://doi.org/10.1016/j.socnet.2007.04.006
https://doi.org/10.1016/j.socnet.2007.04.006
Liben-Nowell, D., & Kleinberg, J. (2007). The link-prediction problem for social networks. *Journal of the American Society for Information Science and Technology*, *58*(7), 1019–1031. https://doi.org/10.1002/asi.20591

Massol, F., Macke, E., Callens, M., & Decaestecker, E. (2021). A methodological framework to analyse determinants of host–microbiota networks, with an application to the relationships between *Daphnia magna'* s gut microbiota and bacterioplankton. *Journal of Animal Ecology*, *90*(1), 102–119. https://doi.org/10.1111/1365-2656.13297

Newman, M. E. J. (2001). Scientific collaboration networks. II. Shortest paths, weighted networks, and centrality. *Physical Review E*, *64*(1), 016132. https://doi.org/10.1103/PhysRevE.64.016132

Ng, A. Y., Jordan, M. I., & Weiss, Y. (2002). *On Spectral Clustering: Analysis and an algorithm*. 849–856.

Page, L., Brin, S., Motwani, R., Winograd, T., & others. (1999). *The pagerank citation ranking: Bringing order to the web*.

Poisot, T. (2023). Guidelines for the prediction of species interactions through binary classification. *Methods in Ecology and Evolution*, *14*(5), 1333–1345. https://doi.org/10.1111/2041-210X.14071

Van Kleunen, L., Dee, L. E., Wootton, K. L., Massol, F., & Clauset, A. (2024). *Predicting missing links in food webs using stacked models and species traits*. Ecology. https://doi.org/10.1101/2024.11.22.624890

Von Luxburg, U. (2007). A tutorial on spectral clustering. *Statistics and Computing*, *17*(4), 395–416. https://doi.org/10.1007/s11222-007-9033-z

White, S., & Smyth, P. (2005). A Spectral Clustering Approach To Finding Communities in Graphs. *Proceedings of the 2005 SIAM International Conference on Data Mining*, 274–285. https://doi.org/10.1137/1.9781611972757.25

Yang, K.-C., Aronson, B., & Ahn, Y.-Y. (2020). BiRank: Fast and Flexible Ranking on Bipartite Networks with R and Python. *Journal of Open Source Software*, *5*(51), 2315. https://doi.org/10.21105/joss.02315

Zhu, M., & Ghodsi, A. (2006). Automatic dimensionality selection from the scree plot via the use of profile likelihood. *Computational Statistics & Data Analysis*, *51*(2), 918–930. https://doi.org/10.1016/j.csda.2005.09.010
